# Aneuploidy-induced proteostasis disruption impairs mitochondrial functions and mediates aggregation of mitochondrial precursor proteins through SQSTM1/p62

**DOI:** 10.1101/2024.07.29.605607

**Authors:** Prince Saforo Amponsah, Jan-Eric Bökenkamp, Svenja Lenhard, Christian Behrends, Johannes Martin Herrmann, Markus Räschle, Zuzana Storchová

## Abstract

Aberrant chromosomal content, or aneuploidy, profoundly affects cellular physiology. Even a gain of a single chromosome disrupts proteostasis due to overexpression of numerous proteins. Consequently, cells accumulate SQSTM1/p62-positive cytosolic bodies and show altered proteasomal and lysosomal activity. To elucidate the p62 interaction network in aneuploid cells, we conducted p62 immunoprecipitation and proximity labeling assays followed by mass spectrometry analysis. Our investigation revealed the enrichment of mitochondrial proteins within the cytosolic p62 interactome and proxitome in aneuploid cells, but not in the proxitome spatially confined to autophagosomes. Immunofluorescence microscopy confirmed increased colocalization of p62 with novel interactors and with mitochondrial proteins in polysomic cells. Moreover, we observed mitochondrial defects characterized by increased perinuclear clustering, reduced oxygen consumption, and reduced mitochondrial DNA abundance in polysomic cells. Furthermore, we demonstrate that polysomic cells exhibit reduced import of mitochondrial proteins and accumulation of mitochondrial precursor proteins in the cytosol. Our data suggest that proteotoxic stress induced by chromosome gains leads to the sequestration of mitochondrial precursor proteins into cytosolic p62-bodies and compromises mitochondrial function.

## Introduction

While most healthy human cells maintain a diploid chromosome content, approximately 90 % of solid tumors contain cells with aberrant chromosome numbers, known as aneuploidy (*1*). This is often associated with a high rate of chromosome segregation errors, referred to as chromosomal instability (CIN). The persistent occurrence of chromosome gains and losses results in the generation of novel karyotypic variations (*2*). Analyses of large cancer datasets as well as functional analyses of aneuploid cells elucidated the pivotal role of aneuploidy in tumorigenesis in promotion of tumor growth, metastasis, and the development of drug resistance (*3, 4*).

Yet, aneuploidy comes at a cost. The gain or loss of even a single chromosome is generally incompatible with the viability of human embryos. The few exceptions to this rule, such as trisomy of chromosomes 13, 18, or 21, are often accompanied by a multitude of pathologies. Lab-engineered human constitutively aneuploid cells manifest numerous defects. Cells lacking a chromosome (monosomy) show reduced proliferation, reduced translation and increased DNA damage, and their viability is incompatible with the presence of functional *TP53* (*5*–*7*). Human cells with extra chromosomes (polysomy) also frequently exhibit slower proliferation rates when compared to their diploid counterparts. Moreover, they manifest an array of phenotypic changes, including features of proteotoxic stress, as evidenced by an increased sensitivity to inhibitors targeting protein folding and turnover, elevated replication stress and accumulation of DNA damage and structural rearrangements, activation of the innate immune response and reduced translation (*5, 8*–*13*). Similar effects were also observed in acute response to aneuploidy, immediately after induced chromosome missegregation (*14*–*16*).

The primary cause of the stresses associated with chromosome gains is largely considered to be the increased expression of several hundreds or thousands of surplus proteins encoded on the extra chromosome. These additional proteins overwhelm the machinery dedicated to protein synthesis, folding, and degradation, leading to proteotoxic stress. Indeed, aneuploid cells with supernumerary chromosomes often activate integrated stress response and unfolded protein response, accumulate protein aggregates, and exhibit defects in protein folding as well as increased autophagy and proteasomal activity (*9, 12, 15, 17*). Oxidative stress and increased production of reactive oxygen species have also been observed in both immediate and late responses to chromosome missegregation. Among the most striking features observed in aneuploid cells is the accumulation of cytosolic deposits positive for sequestosome 1 (SQSTM1/p62), an autophagy receptor and ubiquitin-binding protein that sequesters misfolded proteins, protein aggregates and damaged organelles, and targets them for degradation. Cytosolic p62-bodies accumulate in aneuploid cells arising immediately after induced chromosome missegregation as well as in cells with constitutive chromosome gain (*8, 14*–*16, 18*). The causes of p62 accumulation remain only partially understood. In acute response to chromosome missegregation, p62, which is an autophagy substrate, accumulates due to saturated autophagy (*14, 15*). In cells with constitutive chromosome gain, the increased expression of p62 is observed on both transcriptome and proteome levels (*8, 16, 18*), likely in response to the excess or mis-folded proteins, or in response to increased oxidative stress. Taken together, accumulation of cytosolic p62 bodies is integral to the cellular response to aneuploidy, although their function remains unclear.

p62 is a versatile protein with crucial functions in cellular protein homeostasis and signaling pathways. It acts as a receptor in selective autophagy, where it recognizes ubiquitinated cargo and interacts with autophagy-related proteins, such as LC3, to facilitate cargo engulfment into autophagosomes (*19*). Additionally, p62 serves as a scaffold protein, coordinating the assembly of autophagy-related signaling complexes, thereby regulating the initiation and progression of autophagy. The ability to link cargo recognition, autophagosome formation, and signaling makes p62 a key regulator of the autophagic process (*20, 21*). Moreover, due to its ability to recognize and interact with ubiquitin, p62 connects ubiquitinated proteins to the autophagic machinery, facilitating their degradation through selective autophagy (*22*–*24*). p62 also interacts with numerous signaling molecules, including kinases and transcription factors, regulating cellular processes such as oxidative stress response, inflammation, and cell survival (*21, 25*). Dysregulation of p62 has been implicated in various pathological conditions, including neurodegenerative disorders and cancer, highlighting its importance in protein aggregation and clearance (*26*–*28*). Additionally, p62 acts as a key mediator in mitochondrial turnover, as it functions as an adaptor protein for PINK and Parkin mediated mitophagy as well as an interaction partner of the E3 ubiquitin ligase complex KEAP1 and RBX1 in PINK and Parkin independent mitophagy (*29, 30*). Besides that, p62 regulates mitochondrial dynamics by modulating the activity of DRP1, a protein involved in mitochondrial fission (*31*). However, it remains unclear which of the many functions of p62 is required in response to chromosome gain.

Here, we used our previously established model system of human cell lines engineered to carry one or two extra chromosomal copies (trisomy and tetrasomy, hereafter polysomy) to analyze the function of p62 in response to aneuploidy. We found that the abundance of p62 scales with the number of aneuploid chromosomes in model systems as well as in cancer cell lines of the Cancer Cell Line Encyclopedia (CCLE) based on the analysis of multi-omics data retrieved from the Dependency Map (DepMap) data portal (*32*). Immunoprecipitation of p62 interactors and labeling of p62-proximal proteins via APEX2-p62-dependent biotinylation, followed by mass spectrometry, revealed that a large fraction of p62 interactors in ane-uploid cells comprises of mitochondrial proteins. These findings were further corroborated with observed increased colocalization of p62 with mitochondrial markers. We show that the sequestration of mitochondrial proteins into p62-bodies is a response enhanced in aneuploid cells and can be rescued by improving cytosolic protein homeostasis through expression of protein folding factors. Finally, we demonstrate that proteotoxic stress due to gain of a single chromosome impairs mitochondrial precursor protein import, which leads to their sequestration into p62-positive cytosolic bodies.

## Results

### Abundance of p62 in aneuploid cells scales with the amount of extra DNA

We and others have previously shown that aneuploid cells accumulate autophagy and lysosomal markers on both transcriptome and proteome levels (*8, 14*–*16, 18*). Among the most abundant proteins within this category was the selective autophagy receptor p62, which forms highly abundant cytosolic p62-bodies in polysomic cells. To evaluate the composition of the p62-bodies, we used engineered cell lines with extra chromosomes 3, 5, 13 and 21 transferred into pseudodiploid HCT116 cell line via microcell mediated chromosome transfer (Fig. 1a). The karyotypes were confirmed by chromosome painting and sequencing (*8, 33*). The cell lines were labeled according to their karyotype as HCT116 3/3 (trisomy of chromosome 3), 5/3 (trisomy of chromosome 5), 5/4 (tetrasomy of chromosome 5), 13/3 (trisomy of chromosome 13) and 21/3 (trisomy of chromosome 21). Immunoblotting of cell lysates from these five polysomic cell lines, compared to the parental diploid, revealed that the expression of p62 increases with the increased size of the extra chromosome, hence with the increasing number of additional protein-coding genes (Fig. 1b, c). The p62 abundance further increased upon treatment with an inhibitor of autophagosome-lysosome fusion Bafilomycin A1 (BafA1), confirming the existence of normal autophagic flux in the polysomic cells as previously reported (*8, 18*). The size and number of p62 bodies per cell surface area also increase in polysomic cells (Fig. 1d-f). It should be noted that SQSTM1 is located on chromosome 5 and thus increased expression is expected in trisomy and tetrasomy of chromosome 5. However, the p62 abundance increased in all polysomic cells and the number and size of the bodies tightly correlated with the amount of surplus protein coding genes irrespective of the identity of the extra chromosome (compare trisomy of chromosome 5 and trisomy of chromosome 3, Fig. 1g).

**Figure 1.**
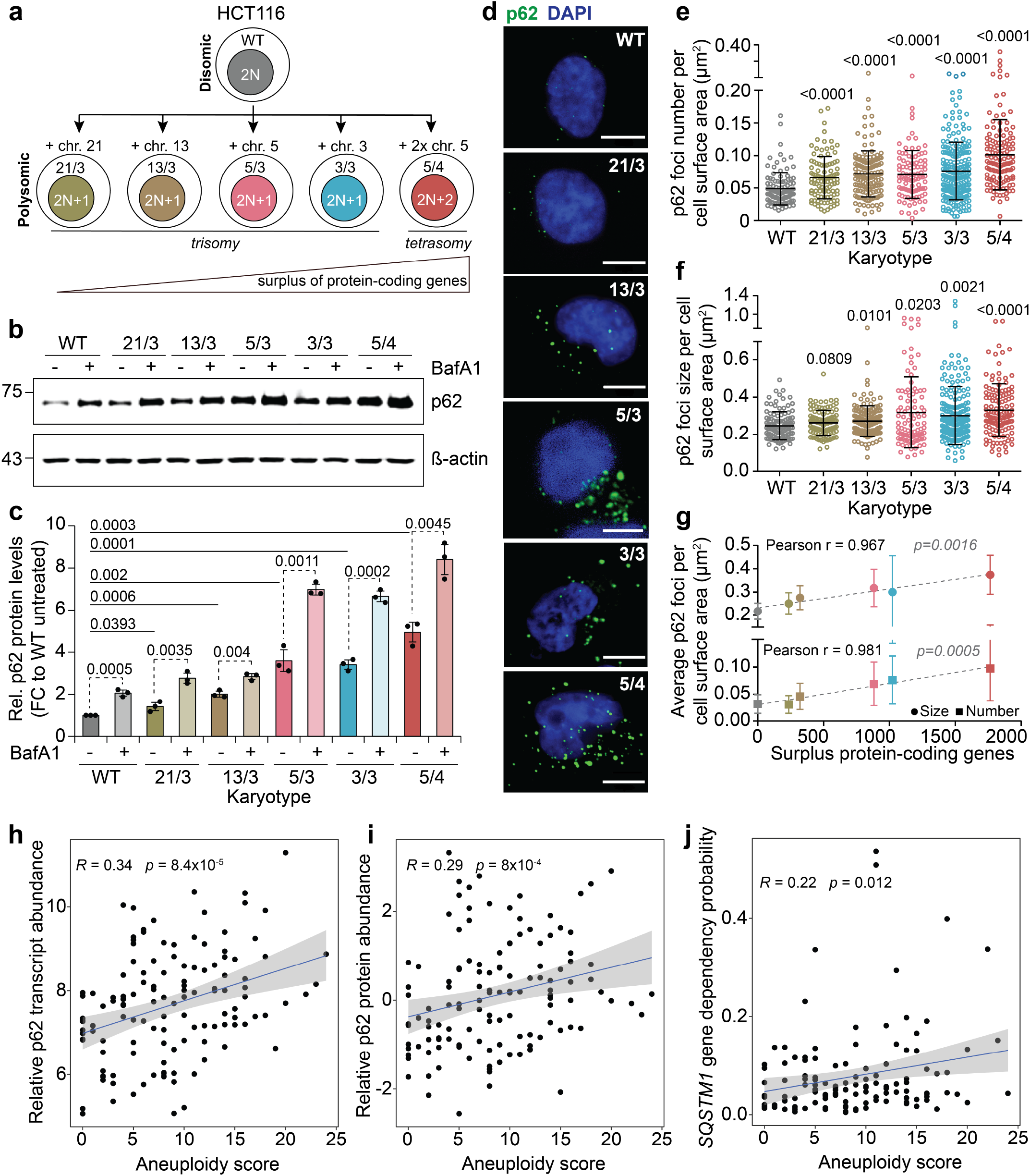
Expression levels and cytosolic deposits of p62 are increased in polysomic cells and correlates with aneuploidy score. **a**, Schematic depiction of used cell lines. **b**, Representative immunoblot of p62 without and with Bafilomycin A1 (BafA1) treatment. ß-actin serves as a loading control. **c**, Quantification of p62 expression levels without and with BafA1 treatment. Data is shown as mean ± s.d. fold change to WT from *n* = 3 independent experiments, and individual replicates are plotted as dots. *P*-values represent two-tailed unpaired Student’s *t*-test. **d**, Representative confocal images of p62 foci immunofluorescence. Scale bar 10 µm. **e, f**, Quantification of p62 foci number and p62 foci size per cell surface area in µm^2^. Individual data for each cell from *n* = 3 independent experiments and means with standard deviation are shown. *P*-values represent non-parametric ANOVA (Kruskal–Wallis statistic for (**e**) 99.54, p *<* 0.0001; for (**f**) 30.37, p *<* 0.0001) followed by Dunn’s multiple comparisons test. **g**, Pearson correlation coefficient of mean ± s.d. p62 foci number and size with the number of surplus protein-coding genes in polysomic cell lines. Two-tailed *P*-values are indicated. **h**, Correlations of relative transcript abundance, **i**, relative protein abundance, and **j**, gene dependency probability, with aneuploidy score of 127 cancer cell lines from the Cancer Cell Line Encyclopedia (CCLE). R represents Spearman’s rank correlation coefficient. *P*-values are two-sided.

The correlation between aneuploidy and p62 abundance is not restricted to model polysomic cell lines. Integration of the genomic (*34, 35*), transcriptomic (*36*), and proteomic (*37*) data obtained for cancer cell lines of the CCLE and stored in the DepMap database (*32*) showed that both the transcript and protein abundance of p62 increases with aneuploidy score, which is defined as the number of chromosome arm-level aberrations (*38*) (Fig. 1h, i). Moreover, the probability of gene dependency for *SQSTM1*, which estimates the reliance of cancer cells on a particular gene, also correlates with the aneuploidy score of cancer cells (Fig. 1j). Together, these data establish that p62 abundance increases with increasing degree of aneuploidy in cancer cells.

### Mitochondrial proteins are predominant interactors of p62 in polysomic cells

To assess the function of p62 in polysomic cells, we decided to identify its interactors. To this end, we performed immunoprecipitation of p62 from whole cell lysates of two polysomic cell lines, HCT116 13/3 and 5/4, and the parental cell line HCT116, followed by mass spectrometry analysis (IP-MS). Since p62 is a substrate for autophagy, we also treated the cells with BafA1 to stabilize p62 levels to enrich the fraction of potential interactors. The data was normalized to a pull down performed in the same lysates using normal IgG as bait (Fig. 2a). In total we found 435-941 proteins enriched with p62 from quadruplicate experiments (Supplementary Data Fig. 1a, b). Principal component analysis showed clustering of quadruplicate measurements across cell lines, treatment conditions and baits (Supplementary Data Fig. 1c). We assessed the statistical significance of differential protein abundance values between p62 pulldowns and the IgG controls to derive a set of candidate interactors of p62 for each karyotype and treatment (Supplementary Table 1, see Methods). For example, we identified NBR1, TAX1BP1, KEAP1 and other well-known interactors of p62 (*39*) in all analyzed cell lines independent of karyotype (Fig. 2b, Extended Data Fig. 1a, b, cluster I). Additionally, we identified candidates exclusive to both polysomic cell lines: the molecular chaperone HSPB1 (HSP27), lysosomal protease CTSZ, mRNA surveillance and ribosome quality control factor PELO, as well as the mitochondrial hydroxyacyl-CoA dehydrogenase trifunctional multienzyme complex subunits HADHA and HADHB (Fig. 2b, Extended Data Fig. 1a, b, cluster II). To validate the interactors, we evaluated their colocalization with p62 by immunofluorescence microscopy. This analysis confirmed colocalization of p62 with the previously identified interactors NBR1, TAX1BP1 and KEAP1 (Fig. 2c, Extended Data Fig. 2c). The newly identified and previously unknown polysomy-specific p62 interactors CTSZ and PELO also colocalized with p62 in polysomic, but rarely in the parental cell line. Importantly, this colocalization was also observed in the 3/3 cell line, which was not subjected to IP-MS, thus confirming the generality of our findings for polysomic cells (Extended Data Fig. 2c).

**Figure 2.**
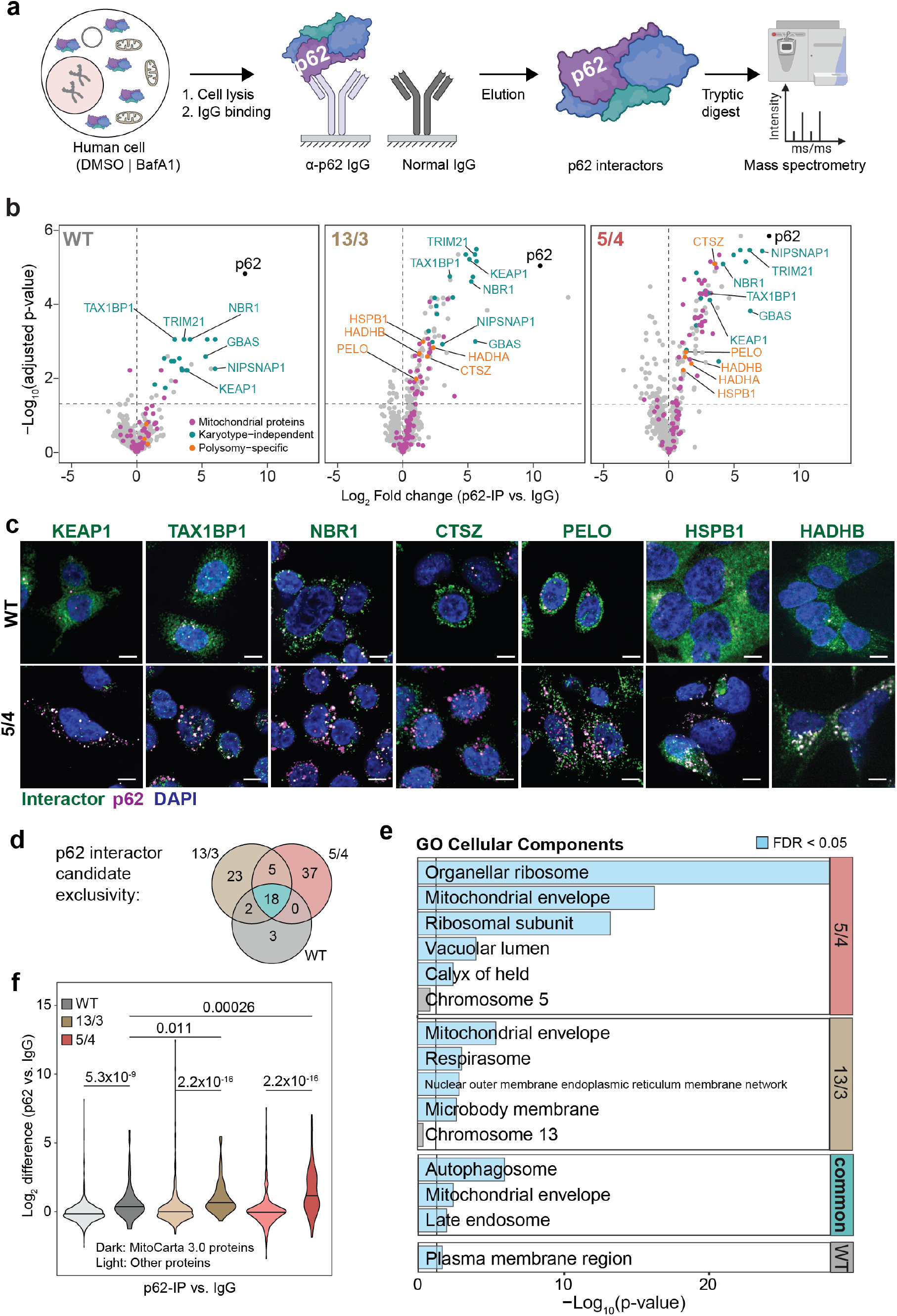
Co-immunoprecipitation followed by label free proteomics reveal mitochondrial proteins are predominant interactors of p62 in polysomic cells. **a**, Schematic depiction of experimental procedure for p62 pull down followed by mass spectrometry. **b**, Volcano plots showing log2-transformed fold change of enriched p62 interactors in WT, 13/3 and 5/4 cells treated with BafA1, highlighting mitochondrial (magenta), karyotype-independent (cyan) and polysomic-specific (orange) interactors. **c**, Representative confocal images of p62 colocalization with selected interactors in WT and 5/4 cells. **d**, Venn diagram showing distribution of enriched p62 interactors between karyotypes when compared to IgG control from all groups in a BafA1 treated cells. **e**, Gene ontology (GO) over-representation analysis of enriched p62 interactors in (**d**). Blue bars represent negative log-transformed *P*-values for GO terms with FDR *<* 0.05. Representatives from clusters of related GO terms are shown (see Methods). **f**, Comparison of fold differences in enrichment of mitochondrial proteins, as defined by the MitoCarta 3.0 inventory, to all other proteins in the measured p62 interactome (**b**). *P*-values are derived from two-sided Wilcoxon’s ranks sum tests.

By overlapping the sets of candidates between the karyotypes we determined common, polysomy-specific as well as karyotype-dependent p62 interactors in both BafA1 treated and untreated samples (Fig. 1b, Extended Data Fig. 1a, Methods). Common hits (18 proteins in BafA1 treated samples, 7 proteins in untreated samples) included p62 interactors that were highly enriched in all cell lines (Fig. 2d, Extended Data Fig. 1d). To further characterize the p62 interactors, we determined whether any cellular compartments (*40, 41*) are significantly over-represented among the candidates in the different karyotypes (Supplementary Table 1). We confirmed that the common p62 interactors are mainly components of the autophagy pathway, which include several mitochondrial proteins (Fig. 2e, Extended Data Fig. 1e). Interestingly, we found no enrichment for proteins encoded on the additional chromosome (Fig. 2e, Extended Data Fig. 1e). Instead, there was a marked over-representation of mitochondrial proteins among the specific interactors identified in polysomic cell lines, but not in the wild type, in both BafA1 treated and untreated samples (Fig. 2e, f, Extended Data Fig. 1e, f). The finding was further corroborated by using the robust MitoCarta3.0 inventory (*42*) of mitochondrial proteins for testing the significance of the enrichment. Interestingly, the specific mitochondrial proteins interacting with p62 differed in the two polysomic cell lines. Since we could not detect any significant association with the individual polysomic chromosomes (chr. 5, or 13), this finding suggests that there are chromosome-specific features that also affect the common phenotypes. Taken together, mitochondrial proteins are the predominant interactors of p62 in cells with additional chromosome.

### p62-dependent cytosolic proximity labeling confirms juxtapo-sition with mitochondrial proteins

To complement the IP-MS experiments, we analyzed p62-proximal proteins by performing APEX2-p62 based proximity biotinylation proteomics (*39*). To this end, we stably expressed *myc-APEX2-SQSTM1* construct in diploid, 13/3 and 5/4 cells by lentiviral transduction (Extended Data Fig. 2a). Transduced cells were then treated with biotin-phenol for 30 min and a pulse (1 min) of 1 mM H_2_O_2_ to induce the biotinylation reaction. Biotinylated proteins were enriched by streptavidin pull down and analyzed by mass spectrometry (Fig. 3a). By this approach, we identified 2321-2874 proteins biotinylated in the proximity of p62 in the cytosol (Supplementary Data Fig. 1d). Principal component analysis showed clustering of quadruplicate measurements across cell lines (Supplementary Data Fig. 1e). To adjust for differences in total protein abundance between the cell lines, we analyzed global protein abundance by separate quantitative proteomics experiments. The fold changes in p62-proximal proteins (hereafter called proxitome) in the polysomic cells relative to the parental diploid were then statistically tested against the corresponding fold changes in global protein abundance. From this analysis, we identified 236 and 520 proteins significantly increased in the p62 proxitome of 13/3 and 5/4 cell lines, respectively (Fig. 3b, Supplementary Table 2). Cellular compartment over-representation analysis showed that mitochondrial proteins are the predominant candidates within the polysomy-specific proxitome of p62 (Fig. 3c, Extended Data Fig. 2b, Supplementary Table 2). The fraction of mitochondrial proteins within the p62 proxitome of the polysomic cells was more than 2-fold higher than their fraction in the total proteome of these cells (Fig. 3d). Additionally, proteins associated with external encapsulating structure organization as well as extracellular matrix were enriched in both polysomic cells, and proteins related to chaperone and chaperonin complexes were enriched in the 5/4 cell line (Fig. 3c). Similarly, as in the IP-MS experiment, we found that the p62 proxitome is not significantly enriched for proteins encoded on the additional chromosomes. Thus, proximity labeling confirms that the p62 interactors are enriched for mitochondrial proteins in cells with extra chromosomes.

**Figure 3.**
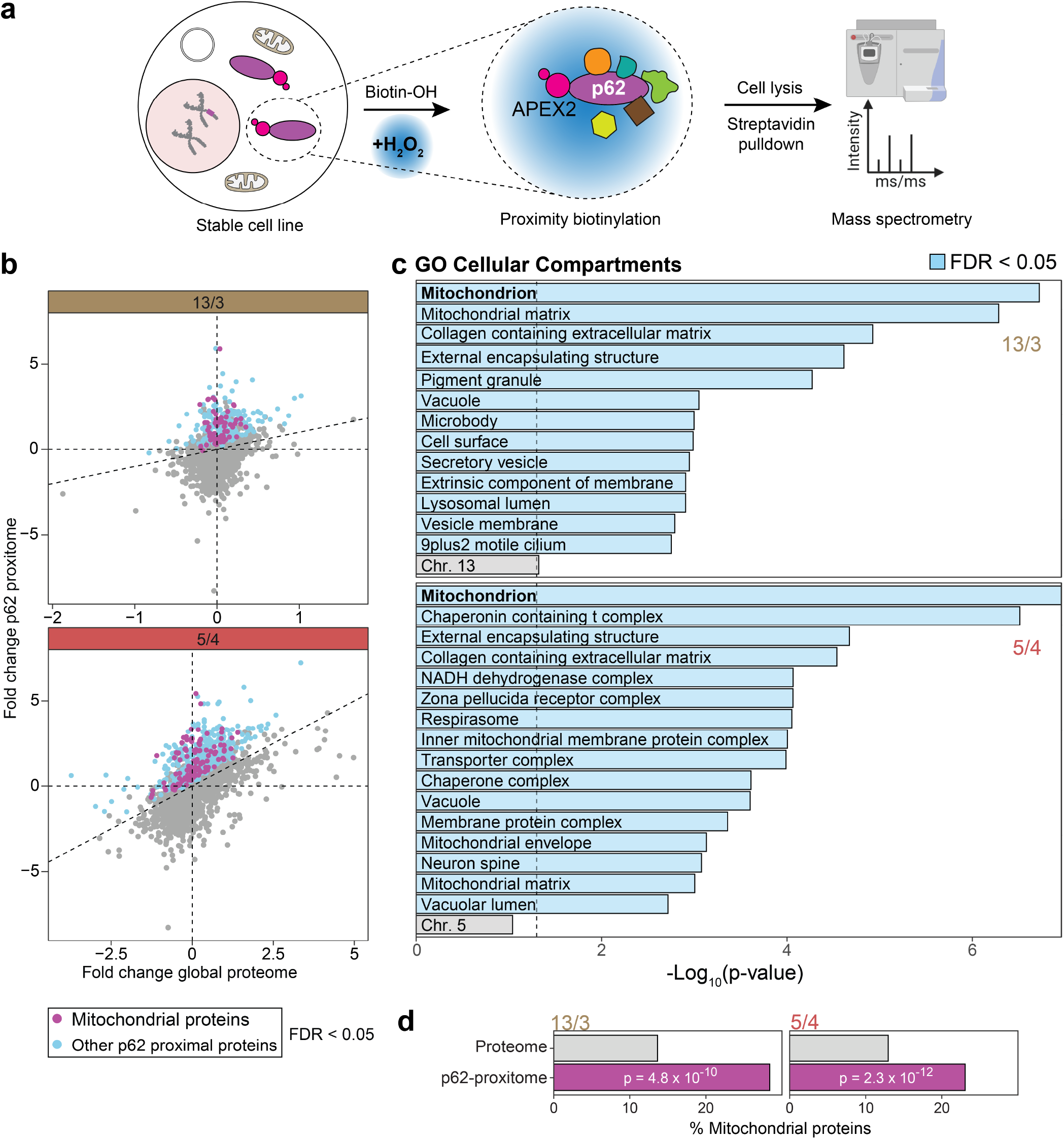
APEX2-based proximity biotinylation of p62 interactors followed by label free proteomics validates enrichment of mitochondrial proteins in polysomic cells. **a**, Schematic depiction of experimental procedure for mass spectrometry-based identification of p62-proximal proteins through APEX2-mediated proximity biotinylation. **b**, Scatter plot of the fold changes in global protein abundance in the 13/3 and 5/4 polysomic cell lines relative to the parental cell line (WT) and the corresponding changes in abundance of cytosolic p62-proximal proteins. Colored dots represent proteins with significantly higher abundance changes of p62-proximal proteins in polysomic cells (FDR *<* 0.05). **c**, Gene ontology (GO) over-representation analysis of the enriched p62-proximal proteins in (**b**). Blue bars represent negative log-transformed *P*-values for GO terms with FDR *<* 0.05. Representatives from clusters of related GO terms are shown (see Methods). **d**, Percentage of mitochondrial proteins in the measured global proteome and the corresponding percentage within the p62 proximal proteome of 13/3 and 5/4 cells. *P*-values represent results of hypergeometric test, evaluating the differences in representation of mitochondrial proteins relative to the measured proteome.

To analyze the p62 proximal mitochondrial proteins in detail, we evaluated their distribution by sub-compartmental localization. This revealed that proteins of the inner membrane (IM) and mitochondrial matrix were enriched in the polysomy-specific p62 proxitome (Extended Data Fig. 2c). Additionally, outer mitochondrial membrane (OM) proteins were significantly present in the p62 proxitome of the 5/4 cells (Extended Data Fig. 2c). By using the TargetP 2.0 prediction tool (*43*), we found a significantly increased enrichment of proteins that are predicted to possess matrix targeting sequences (MTS) in the 13/3 cell line, while there appears to be equal distribution of MTS and non-MTS containing proteins in the 5/4 cell line (Extended Data Fig. 2d). The p62 proxitome in the 5/4 cell line was also significantly enriched with hydrophobic proteins, while no enrichment for subunits of macromolecular complexes was identified (Extended Data Fig. 2e, f). Taken together, our data indicate that p62 bodies in polysomic cells are enriched for inner membrane and mitochondrial matrix proteins with MTS, which is suggestive of impaired mitochondrial import and function.

### Spatially restricted p62-dependent proximity labeling does not show enrichment for mitochondrial proteins in autophagosomes

The autophagy receptor p62 is known for its role in targeting selected cargo to autophagosomes. To elucidate whether the mitochondrial p62-interactome and proxitome in aneuploid cells are channeled through autophagy, we profiled the content of autophagosomes by proximity labeling using APEX2-p62 and limited proteolysis (*39*). Confocal microscopy of the transduced cell lines revealed colocalization of the myc-APEX2-p62 bait and biotin-positive puncta, which in turn colocalized with the lysosomal marker LAMP1 upon BafA1 treatment and proximity labeling (Extended Data Fig. 3a). Immunoblotting confirmed that the APEX2 fusions, biotinylated proteins, and endogenous p62 are partly protected from proteinase K, and enriched for lipidated LC3B-II form (Extended Data Fig. 3b). These results indicate that the APEX2-p62 chimeras are targeted to autophagosomes and biotinylate engulfed proteins, as previously shown (*39*). To perform the analysis, we treated the transduced cells with biotin-phenol (30 min) and subsequent pulsing with H_2_O_2_ (1 min). Clarified lysates were incubated with proteinase K (PK) to digest proteins that were not protected by intact membranes (Fig. 4a).

**Figure 4.**
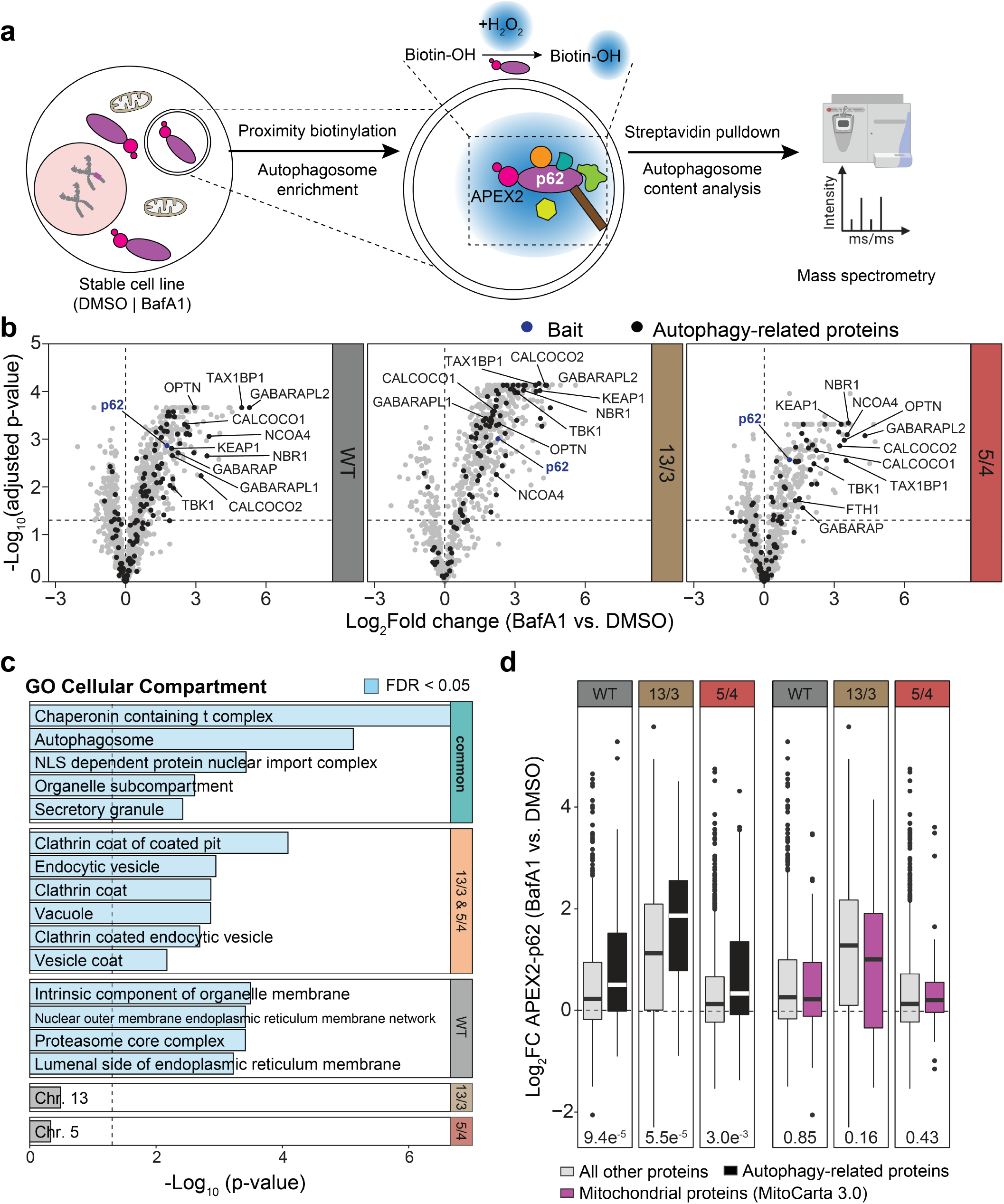
Mitochondrial proteins are not overrepresented p62 cargo candidates in autophagosome lumen. **a**, Schematic depiction of experimental procedure for autophagosome content profiling to identify p62 cargo candidates via APEX2-mediated proximity biotinylation. **b**, Volcano plots showing log2-transformed fold change in BafA1 versus DMSO (vehicle) treated conditions across the used cell lines. Selected known p62 interactors are annotated. **c**, Gene ontology (GO) over-representation analysis of the p62 cargo candidates common to all karyotypes, as well as of additional p62 cargo candidates shared between 13/3 and 5/4 and found in each karyotype separately (Extended Data Fig. 3c). Blue bars represent negative log-transformed *P*-values for GO terms with FDR *<* 0.05. **d**, Boxplots indicating fold changes in abundance of autophagy related proteins (black) and mitochondrial proteins (magenta) compared to all other proteins (grey) enriched among the p62 cargo candidates from (**b**). *P*-values are derived from two-sided Wilcoxon’s ranks sum tests.

MS was conducted after lysis of the enriched autophagosomes and pulldown of the biotinylated proteins using streptavidin agarose, in both BafA1 treated and untreated cells (Fig. 4a). We included a PK resistant control, in which cleared lysates were treated with both PK and RAPIGest to identify proteins not digested by PK. We detected 165 PK-resistant proteins, which were excluded from our analyses (Supplementary Data Fig. 1f). In total, we identified 422 – 1171 p62 proximal proteins enriched in the autophagosome lumen of the cell lines analyzed (Supplementary Data Fig. 1f). Principal component analysis showed clustering of triplicate measurements across cell lines and treatment conditions (Supplementary Data Fig. 1g). First, we asked what proteins are enriched among p62 proximal proteins in the autophagosome lumen of polysomic cells compared to parental cells in normal conditions (no Baf1A treatment). To this end, we tested the fold changes in p62 proxitome in autophagosomal lumen of the polysomic cells normalized to parental diploid against the corresponding fold changes in global protein abundance (Extended Data Fig. 4a) and identified 109 and 184 significantly enriched proteins in the 13/3 and 5/4 cell lines, respectively (Extended Data Fig. 4a, Supplementary Table 3). While this included several mitochondrial proteins, there was no significant over-representation of mitochondrial proteins. Instead, cytosolic ribosomes and ribonucleoproteins were the predominant candidates over-represented within the polysomy-specific autophagosome lumen p62 proxitome (Extended Data Fig. 4b, c, Supplementary Table 3). Next, we assessed the statistical significance of differential protein abundance values between spatially restricted p62 proxitome in BafA1-treated cells, normalized to untreated cells to derive a set of BafA1-stabilized p62 cargo proteins for each karyotype. We observed enrichment for hATG8s (e.g., GABARAP), selective autophagy receptors (e.g., CALCOCO1, CALCOCO2, NBR1), established p62 substrates (e.g., KEAP1), and several known p62 cargo proteins (e.g., TBK1, Fig. 4b, Supplementary Table 3). Autophagy proteins were also significantly over-represented among the identified p62 cargo proteins common to all karyotypes (Fig. 4c). Surprisingly, we found no significant over-representation of mitochondrial proteins in the polysomy-specific p62 cargo in autophagosomal lumen (Fig. 4c, d, Extended Data Fig. 3c). There was also no significant over-representation of proteins encoded by genes on the extra chromosomes. The p62 cargo common to both polysomic cell lines, but not the parental diploid cells, included proteins mainly related to endocytic vesicle formation and release. Together, this suggests that mitochondrial proteins are sequestered into p62-positive cytosolic bodies in polysomic cells, but these proteins are not targeted by p62 into autophagosomes.

### Chromosome gain alters mitochondrial architecture and genome

Since the highly abundant p62 bodies sequester mitochondrial proteins in polysomic cells, we hypothesized that the mitochondria in these cells might be impaired. We reasoned that such an impairment would also lead to colocalization of p62 with TOMM20 (translocase of outer mitochondrial membrane 20), which is often used as an indicator of mitochondrial damage (*44, 45*). Analysis of colocalization of p62 with TOMM20, which was not identified in the IP-MS as an interactor of p62, uncovered low colocalization in HCT116 cells growing in optimal conditions, but a clear increase in polysomic cells (Fig. 5a, b). Quantification of the Pearson correlation coefficient revealed that the colocalization strongly correlates with the number of surplus protein coding genes (Fig. 5c). The immunofluorescence of TOMM20 also showed that the mitochondrial architecture is altered in polysomic cells compared to the parental cell line, as the fraction of cells with perinuclearly clustered mitochondrial network, which is frequently observed upon stress conditions, such as hypoxia, and is mediated by p62 (*46*), was significantly increased in polysomic cells (Fig. 5d, e). Due to the accumulation of perinuclear mitochondria, we hypothesized that most of the mitochondria might be depolarized. To this end, we measured mitochondrial mass and membrane potential by staining the cells with MitoTracker Green FM and DeepRed FM dyes, respectively, and analyzed fluorescence intensity by flow cytometry in the parental diploids, 21/3, 13/3 and 5/4 cells. Surprisingly, the membrane potential, measured as a ratio of MitoTracker DeepRed to MitoTracker Green signal, was slightly higher in polysomic compared to parental cells, while the mitochondrial mass was unchanged (Extended Data Fig. 5a-d). Thus, the gain of a chromosome impairs mitochondria in human cells, but without affecting their membrane potential.

**Figure 5.**
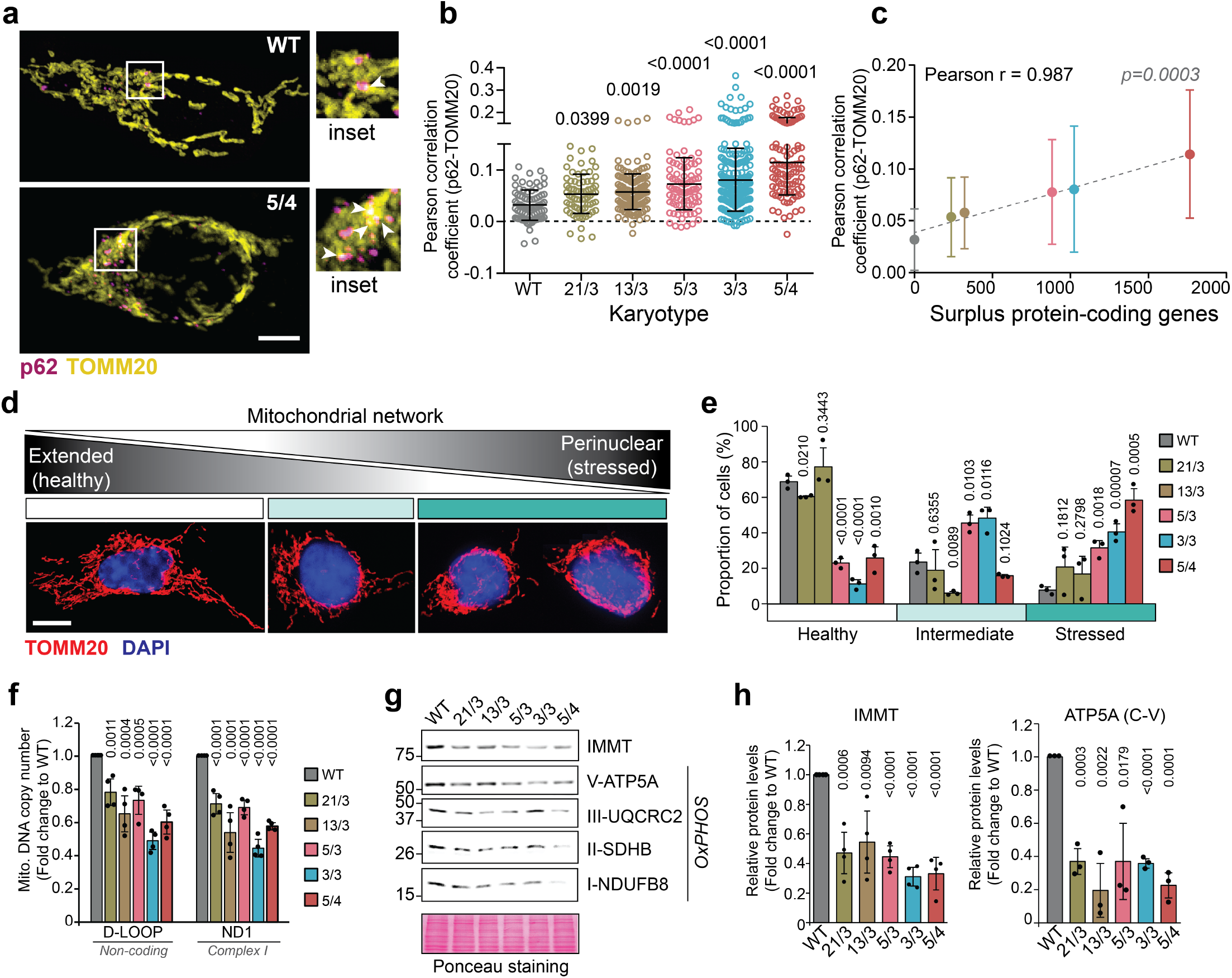
Polysomic cells exhibit increased mitochondrial defects. **a**, Representative confocal images of p62 (magenta) and mitochondrial protein TOMM20 (yellow) colocalization in parental control (WT) and a polysomic (5/4) cell line. Magnified inserts are shown. White arrow heads indicate p62-TOMM20 colocalization. Scale bar 10 µm. **b**, Quantification of the Pearson correlation coefficient for p62-TOMM20 colocalization in WT, 21/3, 13/3, 5/3, 3/3 and 5/4 cells. Values of individual cells from *n* = 3 independent experiments and means with s.d. are shown. *P*-values represent one-way ANOVA followed by Sidak’s multiple comparisons test. **c**, Pearson correlation of p62-TOMM20 colocalization with the number of surplus protein-coding genes. Two-tailed *P*-value is indicated. **d**, Representative confocal images of the different mitochondrial architecture observed in the cells. **e**, Quantification of the cell fractions with each mitochondrial form, classified as healthy (extended network), intermediate (minimally extended network) and stressed (perinuclearly clustered network) as shown in (**d**). Histogram represents average of *n* = 3 independent experiments and individual replicates are shown as dots. *P*-values represent two-tailed unpaired Student’s *t*-test. **f**, Relative mitochondrial DNA (mt-DNA) copy number determined by qPCR of the non-coding mt-DNA D-Loop and respiratory chain complex 1 subunit ND1, normalized to the nuclear ß_2_-microglobulin gene. Data is shown as mean ± s.d. fold change to WT from *n* = 5 independent experiments, and individual replicates are shown as dots. *P*-values represent two-tailed unpaired Student’s *t*-test. **g**, Representative immunoblot and **h**, quantification of expression levels of selected mitochondrial proteins. Ponceau staining is used as loading control. Data is shown as mean ± s.d. fold change to WT from at least *n* = 3 independent experiments, and individual replicates are shown as dot plots. *P*-values represent two-tailed unpaired Student’s *t*-test.

Next, we analyzed the impact of polysomy on the mitochondrial genome. We quantified the mitochondrial DNA copy number (i.e., mtDNA-CN) in polysomic cells in comparison to the parental diploid through quantitative PCR, using primers against the non-coding D-loop and coding sequences of respiratory chain complex or oxidative phosphorylation (OxPHOS) proteins. This analysis revealed that the mtDNA-CN in polysomic cells is reduced to approximately 60 - 80 % of the parental cell line (Fig. 5f, Extended Data Fig. 5e). Given that the mitochondrial mass is unaffected in polysomic cells (Extended Data Fig. 5d), this observation suggests perturbed mitochondrial genome maintenance in response to chromosome gain. Moreover, immunoblotting of selected nuclear encoded OxPHOS proteins, as well as the inner mitochondrial membrane protein IMMT, which we identified in the p62 IP-MS, revealed more than 20 % abundance reduction in polysomic cells (Fig. 5g, h, Extended Data Fig. 5f). In summary, our data suggest that chromosome gain affects mitochondrial ultrastructure and genome maintenance and decreases levels of nuclear encoded respiratory chain components.

### Chromosome gain modifies mitochondrial functions

Having established the enrichment of p62 interactors for mitochondrial proteins in polysomic cells, we focused on further characterization of the nature of the mitochondrial defects using the HCT116 5/4 cell line, where the changes were most prominent. Firstly, we compared the extent of p62 colocalization with mitochondria in the polysomic cells to diploid parental cells where we induced mitochondrial damage using the membrane potential uncoupler CCCP (carbonyl cyanide chlorophenylhydrazone). Staining of mitochondria with MitoTracker Red CMXRos, a chemical dye whose accumulation in mitochondria is independent of membrane potential, revealed two-fold increase of p62-mitochondria colocalization in CCCP-treated compared to untreated parental cells, and equivalent to levels observed in untreated polysomic cells. In contrast, there was no difference in the Pearson correlation coefficient of polysomic cells in the absence and presence of CCCP (Extended Data Fig. 6a), suggesting that a large fraction of mitochondria was defective in polysomic cells even under optimal conditions. Secondly, we isolated mitochondria from polysomic and diploid parental cells and measured their respiration capacity by determining oxygen consumption rate in an oxychamber using NADH as substrate (*47*) normalized to total amount of mitochondrial proteins. Additionally, we included a control reaction in which isolated mitochondria from parental cells were treated with 10 mM CCCP for 15 min prior to oxygen consumption measurement. Polysomic cells showed a significant reduction in oxygen consumption to approximately 50 % of the parental cells and synonymous to CCCP-treated mitochondrial from parental cells (Extended Data Fig. 6b, c). The mitochondrial network in cells undergo constant remodeling through coordinated fusion and fission reactions. Therefore, we evaluated by Western blots the abundance of DRP1 (Dynamin related protein 1), as well as processing of OPA1 (optic atrophy 1), both of which are markers for mitochondrial fission and fusion, from whole cell lysates obtained from the cells. We observed an increased abundance of the processed short form of OPA1 (S - OPA1) in comparison to the total OPA1 levels in the polysomic cell line, as well as increased DRP1 levels. This observation resembled that of parental cells treated with CCCP, which also induces mitochondrial fission, indicating that mitochondrial fission is increased in polysomic cells (Extended Data Fig. 6d, e). These observations suggest that mitochondrial homeostasis is significantly altered in response to chromosome gain.

### Cytosolic proteostasis is critical for formation of p62 bodies and mitochondrial function

Polysomic cells express high levels of p62 and therefore we considered the possibility that the overexpression of p62 alone is sufficient to increase mitochondrial damage in otherwise healthy cells. To this end, we transiently overexpressed EGFP-p62 under the control of a strong CMV promotor (*48*) in the diploid parental cell line HCT116. As a control, we overexpressed the EGFP construct only. While overexpression of EGFP-p62 leads to a strong accumulation of p62-positive bodies in the cytosol compared to the control cells overexpressing EGFP only, there was no increase in colocalization with TOMM20 (Fig. 6a-e). Thus, the increased amount of p62 alone is not the cause of the mitochondrial damage in polysomic cells.

**Figure 6.**
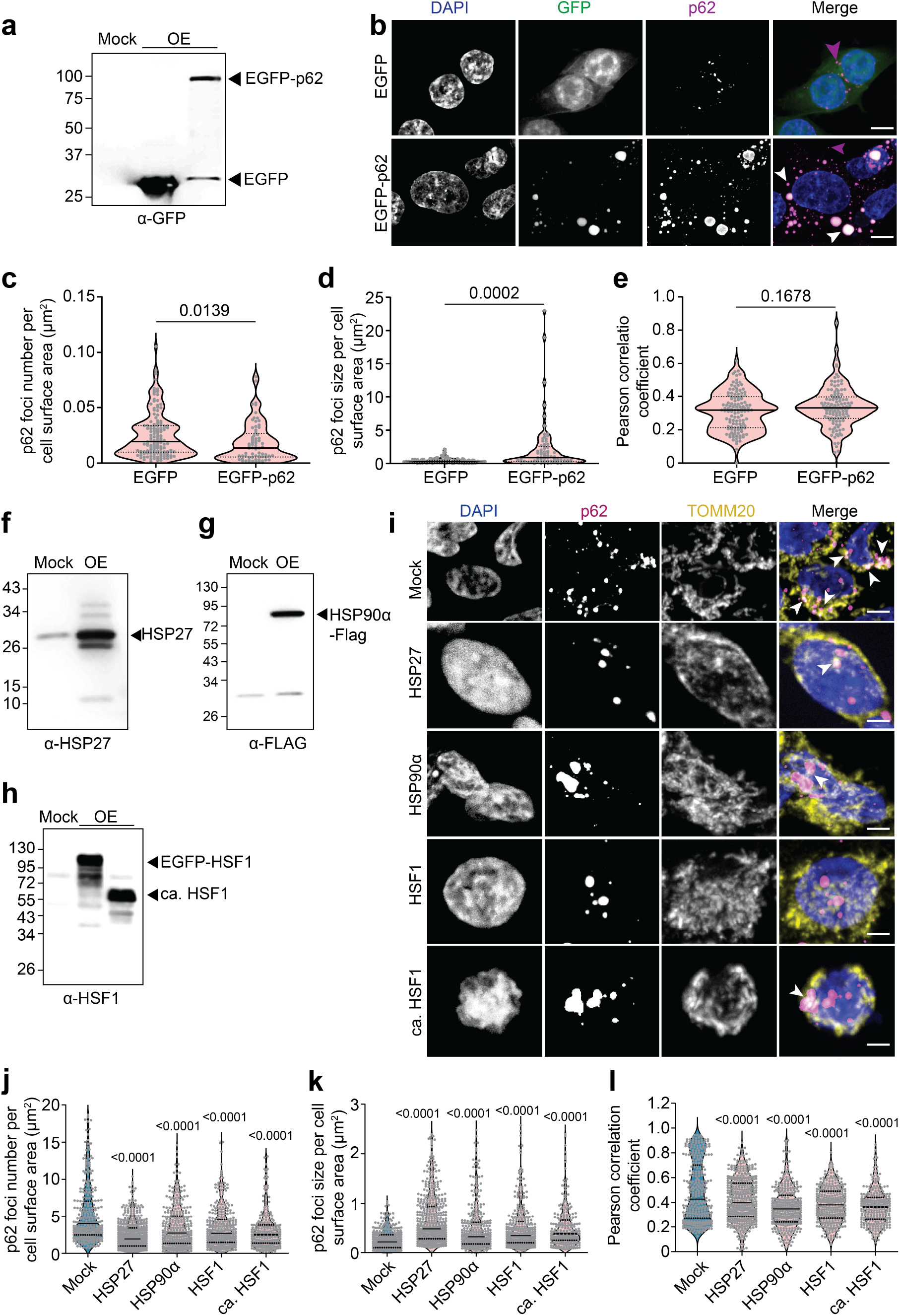
Disruptions in cytosolic proteostasis underlie p62 deposits accumulation and mitochondrial defects, which is ameliorated by transient overexpression of protein folding factors. **a**, Representative immunoblot showing EGFP and EGFP-p62 overexpression. **b**, Confocal images of p62 and EGFP foci visualization in parental control (WT) cells transiently transfected with EGFP and EGFP-p62 plasmids. Scale bar 10 µm. **c**, Quantifications of p62 foci number per cell surface area in µm^2^, **d**, p62 foci size per cell surface area in µm^2^, and **e**, Pearson correlation coefficient for p62-TOMM20 colocalization in respective samples. Violin plots indicate mean and distribution of values, while dots represent individual cells from *n* = 3 independent experiments. *P*-values represent two-tailed unpaired *t*-test with Welch’s correction. **f**, Representative immunoblot showing overexpression of HSP27 **g**, HSP90α-Flag **h**, EGFP-HSF1 and constitutively active HSF1 in 3/3 cell line. **i**, Representative confocal images of p62 foci and TOMM20 visualization in the cells from (**f-h**). Scale bar 10 µm. **j**, Quantification of p62 foci number and **k**, p62 foci size per cell surface area in µm^2^ in overexpressing samples from (**i**). Violin plots indicate mean and distribution of data while dot plots represent individual cells from *n* = 3 independent experiments. *P*-values represent non-parametric ANOVA (Kruskal–Wallis statistic for (**j**) 145.5, p *<* 0.0001; for (**k**) 197.9, p *<* 0.0001) followed by Dunn’s multiple comparisons test. **l**, Pearson correlation coefficient for p62-TOMM20 colocalization in respective samples from (**i**). Violin plots indicate mean and distribution of data while dot plots represent individual cells from *n* =3 independent experiments. *P*-values represent one-way ANOVA followed by Sidak’s multiple comparisons test.

We next hypothesized that impaired cytosolic protein homeostasis due to protein folding defects typical for cells with extra chromosomes (*9, 49, 50*) caused the observed mitochondrial defects in polysomic cells. To test this hypothesis, we sought to reduce the cytosolic proteotoxic stress in polysomic cells by transient overexpression of heat shock protein 27 (HSP27), a small protein chaperone and an antioxidant with a role in stress resistance; alpha subunit of HSP90 (HSP90α), a molecular chaperone required for the maturation and structural maintenance of many proteins, and the wild type and constitutively active variant of the heat shock transcription factor 1 - HSF1 and ca-HSF1, respectively, the transcriptional factor and master regulator of heat shock response. As a control, we used mock transfected 3/3 and 5/4 cells. Overexpression of these constructs were confirmed by Western blot (Fig. 6f-h, Extended Data Fig. 7a). In all instances, we observed decreased numbers of p62-positive foci, but increased foci sizes (Fig. 6i-k, Extended data Fig. 7b, c, d). Pearson correlation coefficient of p62-TOMM20 colocalization revealed significantly reduced colocalization in the 3/3 cells overexpressing the constructs (Fig. 6l), while the 5/4 cells showed no rescue (Extended Data Fig. 7e). We conclude that the proteotoxic stress and mitochondrial defect can be rescued by overexpression of cytosolic chaperones, although only in some polysomic cells. High numbers of extra protein coding genes, and therefore high proteotoxic stress, as in the 5/4 cell line, could not be fully rescued by transient overexpression. Our data suggests that impaired protein folding and chronic overexpression of extra proteins in polysomic cells facilitate formation of cytosolic p62-positive bodies that sequester mitochondrial proteins, thus impairing mitochondrial functions.

### Mitochondrial protein import is diminished in cells with extra chromosomes

Cytosolic protein aggregates can impede mitochondrial biogenesis and function (*51*–*54*). Our data suggests that enrichment of mitochondrial proteins among the p62 interactors in polysomic cells arises due to the accumulation of unimported mitochondrial precursor proteins in cytosolic aggregates, thus altering mitochondrial protein composition and function. To test this possibility, we analyzed the expression of three selected mitochondrial proteins that were identified among p62 interactors through the IP-MS, HADHA, HADHB, and MRPL45, by Western blot in whole cell lysates of the five different polysomic cell lines and parental diploid. Strikingly, we observed increased accumulation of the precursor forms of each of the proteins in whole cell lysates of polysomic cells (Fig. 7a). This indicates that mitochondria of polysomic cells exhibit import defects, despite their normal membrane potential (Extended Data Fig. 5a-c). To validate this observation by an orthogonal approach, we isolated mitochondria from the cells and performed kinetic import of *in-silico* synthesized mitochondria substrates su9-DHFR and human SOD2 (Extended Data Fig. 8a, (*55*)). The mitochondria enrichment was comparable in all cell lines, as confirmed by Western blot (Extended Data Fig. 8b). Kinetic import analysis revealed a delayed import of both su9-DHFR and human SOD2 into isolated mitochondria from polysomic cells (Fig. 7b-e, Extended Data Fig. 8c, d). We propose that cytosolic proteotoxic stress due to defective protein folding leads to mild, but chronic reduction of mitochondrial protein import and impaired mitochondrial function in aneuploid cells. This results in the accumulation of mitochondrial precursor proteins in the cytosol, where they get sequestered into p62-positive bodies (Fig. 7f).

**Figure 7.**
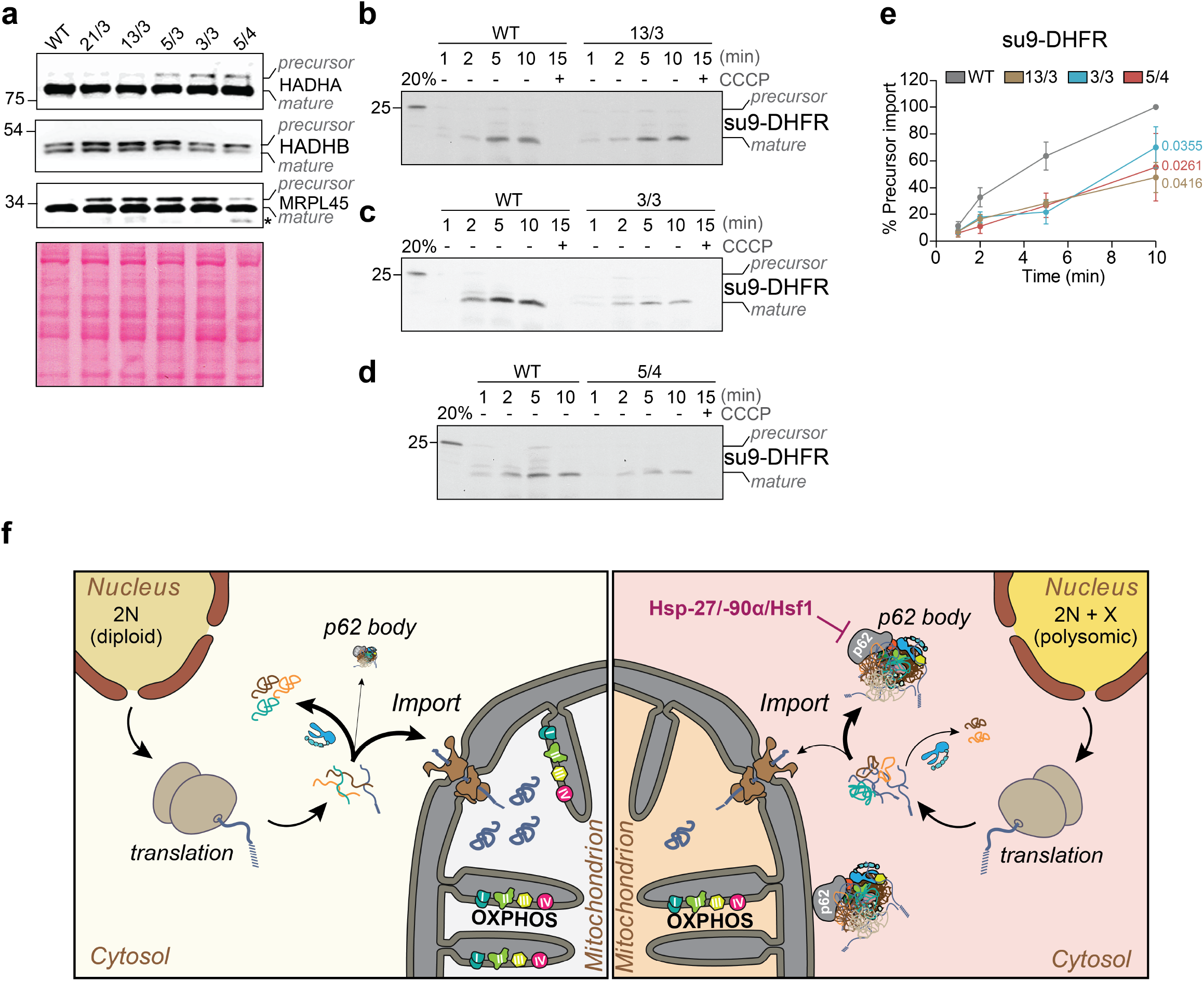
Chromosome gain impairs mitochondrial precursor protein import. **a**, Immunoblot of selected mitochondrial proteins showing precursor and mature forms. Ponceau staining is used as loading control. *Degradation product. **b-d**, Representative images showing import kinetic of *in-silico* synthesized ^35^S-methonine-labeled su9-DHFR into mitochondria isolated from WT, 13/3, 3/3 and 5/4 cells. The su9-DHFR precursor is processed upon reaching the matrix (i.e., mature form). CCCP, which depletes mitochondrial membrane potential thereby preventing import, was used as a negative control. 20 % of the synthesized substrate (precursor) was loaded for comparison. All samples were resolved by SDS-PAGE and visualized by autoradiography. **e**, Quantification of the import kinetic from (**b-d**). Data represents mean ± s.e.m. of *n* = 3 independent import assays. *P*-values represent one-tailed paired *t*-test. **f**, Proposed model of aneuploidy-induced disruptions of cellular proteostasis and their effect on mitochondria. The gain of extra chromosome leads to translation of excess proteins, which overwhelm the protein folding machinery. Aberrant cytosolic proteostasis maintenance arising from defective protein folding impairs mitochondrial function and delays protein import, leading to the accumulation of mitochondrial precursor proteins in the cytosol. The polysomy-induced accumulation of cytosolic p62 bodies (right) results in the sequestration of misfolded proteins and unimported mitochondrial precursor proteins. The p62-positive bodies do not only reside in the cytosol, but also associate with the mitochondrial outer membrane. Overexpression of protein folding factors decreases the formation of p62-positive bodies and their association with mitochondria.

## Discussion

Protein homeostasis is crucial for cellular and organismal functions. Numerical chromosomal aneuploidy, which is associated with various pathologies, alters the abundance of the proteins encoded on affected chromosomes. The mild, but chronic overexpression of hundreds of unneeded proteins in cells with extra chromosomes alters proteome landscape of the cell and triggers aneuploidy-associated stress response. As a consequence, human cells with additional chromosomes suffer from proteotoxic stress, as documented by increased sensitivity to chaperone inhibitors, accumulation of ubiquitin- and p62-positive cytosolic bodies, and increased lysosomal stress (*8, 9, 14*–*17*). By analyzing the phenotypes of five cell lines with different extra chromosomes, we show that the number and size of p62-positive cytosolic bodies tightly correlate with the number of extra protein-coding genes in polysomic cells (Fig. 1a-g). Moreover, the degree of aneuploidy quantified as aneuploidy score in cancer datasets positively correlates with p62 abundance and p62 essentiality (Fig. 1h-j). Thus, p62 expression scales with the protein imbalance caused by chromosome number changes. The reasons for increased p62 abundance in aneuploid cells remain unclear. Acute aneuploidy after chromosome missegregation may lead to p62 accumulation through transient lysosomal stress and autophagy overburdening (*14, 15*). In contrast, chronic aneuploidy does not impair autophagy flux and triggers increased transcription of the SQSTM1 gene in both model cell lines and in cancer datasets ((*8, 18*), Fig. 1). The multifaceted protein p62 has a complex role in maintenance of protein homeostasis, and response to various stresses (*56*). Understanding what triggers accumulation of p62 in response to aneuploidy will be crucial to elucidate its link to cancer.

We hypothesized that analysis of the protein content of the cytosolic p62 bodies in aneuploid cells will provide an insight into the changes caused by the proteotoxic stress in these cells. Proteomic analysis of p62 interactors performed by two complementary approaches revealed a marked complexity of these bodies. Besides previously known interactors, we found and validated several new, aneuploidy specific interactors, and a strong enrichment for mitochondrial proteins (Fig. 2, 3, Extended Data Fig. 1, 2). Remarkably, we did not observe any significant enrichment for proteins encoded on the extra chromosomes, suggesting that p62 does not specifically sequester these supernumerary proteins. Another aspect is the lack of an enrichment of mitochondrial proteins in the spatially restricted p62-proxitome of autophagosomes. This suggests that autophagosome-dependent recycling of damaged mitochondria and mitochondrial proteins in polysomic cells is not mediated via the p62 receptor. In future, profiling of autophagosomes via proximity labeling approaches with other autophagy cargo carriers (*39*) as well as comprehensive analyses of the fate of unimported mitochondrial precursor proteins and damaged mitochondria in polysomic cells will provide further insights about changes in mitochondrial homeostasis in response to aneuploidy.

The p62 bodies often localized near mitochondrial surface in polysomic cells (Fig. 5a-c), a feature which correlates with mitochondrial damage, but not necessarily with mitophagy (*57*). Indeed, cells with extra chromosomes contain stressed, less functional mi-tochondria, and impaired mitochondrial metabolism, reduced levels of mitochondrial proteins and mtDNA (Fig. 5, Extended Data Fig. 5e, f, 6). Aneuploidy has previously been linked to metabolic stress due to increased ROS levels in mitochondria and altered mitochondrial metabolism (*58, 59*). In Drosophila, saturated autophagy in aneuploid cells causes accumulation of dysfunctional mitochondria, increased ROS production and subsequent cellular senescence (*60*). Mitochondrial dysfunction is also frequent in Down syndrome, which is caused by trisomy of chromosome 21. Here, downregulation of nuclear encoded mitochondrial genes, reduced efficiency of energy production, reduced oxygen consumption, and altered mitochondrial morphology were observed (*61*); these features we also observed in engineered polysomic cells regardless the identity of the extra chromosome. While increased expression of genes encoded on chromosome 21, such as the DYRKA1 kinase (*62*), likely contributes to mitochondrial phenotypes, our results suggest that the cytosolic proteotoxic stress caused by production of extra proteins may be another factor causing mitochondrial dysfunction.

What is causing the mitochondrial defects in polysomic cells? Mitochondria contain their own DNA, but 99 % of mitochondrial proteins are encoded in the nucleus, synthesized in the cytosol, and imported into mitochondria by dedicated transport pathways (*63*). Strikingly, there was increased accumulation of mitochondrial precursor proteins in the cytosol of polysomic cells and the import into mitochondria was delayed in polysomic cells (Fig. 7a-e). Mitochondrial inner membrane-localized proteins and mitoribosomal proteins, which are usually hydrophobic in nature, pose severe challenges to cytosolic proteostasis when their import into mitochondria is impaired. Their sequestration into aggregates acts as a protective measure to minimize their negative effect (*64*). We found that these proteins were particularly enriched in the p62-positive bodies in aneuploid cells (Fig. 2, 3). We hypothesize that the impaired protein folding and imbalanced protein homeostasis due to aneuploidy induces protein aggregation (*8, 15, 17, 49, 50, 65*) into cytosolic p62-positive bodies, which sequester unimported mitochondrial proteins. The reduced import of essential mitochondrial proteins into mitochondria impairs mitochondrial health (Fig. 7f). Our findings demonstrate that cytosolic proteotoxic stress in cells with extra chromosome is sufficient to impair mitochondrial functions in human cells.

Recent years have brought progress in our understanding of the intimate link between mitochondrial function and cytosolic protein homeostasis (*66, 67*). Cytosolic protein folding and quality control mechanisms ensure that nuclear encoded proteins are correctly targeted to their mitochondrial destinations and any disruption of this process exacerbates their misfolding and aggregation. Disruptions in cytosolic proteostasis can affect mitochondrial morphology and distribution due to disruption of mitochondrial dynamics. Here we show that aneuploidy, an omnipresent feature of cancer cells, affects protein import into mitochondria and triggers sequestration of mitochondrial proteins into cytosolic aggregates. Our work sheds new insight into consequences of aneuploidy and provide novel, physiological relevant model system to study the link between cytosolic protein imbalance and mitochondrial function.

## Acknowledgments

We thank Ulrich Hartl, Len Neckers, Bianca Brundel and Anne Simonson for the plasmids. We thank Osmar Velazquez, Anna Myronova, Olha Kurpa, Devi Murugan, Caroline Erler, Celina Hirschel-mann and Lena Johann for their help with experiments. This project was supported by grants from the Deutsche Forschungsgemeinschaft (GRK2737-STRESSistance to ZS and JMH, and Walter Benjamin Programme Award AM 703/1 to PSA), and the Landesforschungsinitiative Rheinland-Pfalz BioComp (to ZS and JMH). PSA is additionally supported by the TU Nachwuchsring Kaiserslautern and an Add-on fellowship from the Joachim Herz Stiftung.

## Author Contributions

PSA performed the biochemical, proteomic and cell biology experiments. PSA and MR carried out the mass spectrometry analysis; JEB and MR performed the analysis of the proteomics data, and JEB performed the analysis of CCLE data. SL carried out the protein import experiments. ZS, PSA and JMH supervised the study. PSA, JEB and ZS wrote the manuscript, all authors critically reviewed and commented on the manuscript.

## Online methods

### Cell lines and culture conditions

Polysomic cell lines were derived from the near-diploid human colon carcinoma cell line HCT116 (ATCC No. CCL-247) expressing H2B-GFP by microcell-mediated chromosome transfer as previously described (*8, 33*). Parental HCT116 (WT), HCT116 21/3 (trisomy of chromosome 21), 13/3 (trisomy of chromosome 13), 5/3 (trisomy of chromosome 5), 3/3 (trisomy of chromosome 3), and 5/4 (tetrasomy of chromosome 5) were cultured in Dulbecco’s Modified Eagle Medium (DMEM; Gibco) supplemented with 10 % Fetal Bovine Serum and 5 % Penicillin/Streptomycin and incubated at 37 °C with 5 % CO_2_ atmosphere. All cell lines were used up to a maximum of five passages before replacement with fresh ones and tested regularly for mycoplasma contamination using MycoStrip tests (InvivoGen). Bafilomycin A1 was applied at 100 nm for 6 h, CCCP at 10 µM for 6h.

### Transient cell line transfection

Transient expression of the following plasmids Hsp90α, Hsp27, Hsf1, ca. Hsf1, pEGFP-N3, EGFP-p62 were performed using Lipofectamine 2000 (Thermo Fisher Scientific) according to the manufacturer’s instructions. Transfection medium was replaced 6 h post post-transfection and transfected cells were analyzed 48 h post-transfection. A list of plasmids used is provided in Supplementary Table 4.

### Stable cell line transfection

HCT116 WT, 13/3 and 5/4 cell lines stably expressing myc-APEX2-p62 were generated by lentiviral transduction. Briefly, HEK 293T cells were transiently transfected with the packaging plasmids pHDM-Hgpm2, pHDM-Tat1b, pHDM-VSV-G, and pRC-CMV-Rev1b (*68*) and the expression plasmids using Lipofectamine 2000 (Thermo Fisher Scientific) according to the manufacturer’s instructions. Fourty-eight hours post-transfection recipient cells were infected with the viral supernatant in the presence of 8 µg/ml polybrene according to the manufacturer’s instructions (Sigma-Aldrich). myc-APEX2-p62 transduced cells were selected in cell culture media supplemented with 2 µg/ml puromycin (Thermo Fisher Scientific) A list of plasmids used is provided in Supplementary Table 4.

### Analysis of p62 interactome by co-immunoprecipitation-based quantitative proteomics

For immunoprecipitation, cell lysates were prepared by lysis in buffer [150 mM NaCl, 50 mM Tris-HCl (pH 7.5), 5 % Glycerol, 1 % IGPAL-CA-630, 1 mM MgCl_2_] supplemented with 1 mM PMSF and 1X Protease inhibitior cocktail (Merck). Samples were incubated on an overhead rotator for 20 min and centrifuged at 500 g for 10 min at 4 °C to remove cell debris. Total protein concentration was determined in the supernatant fraction by Bradford assay. SQSTM1 antibody (Santa Cruz, sc-28359) and Normal mouse IgG (Santa Cruz, sc-2025) were added separately to 1 mg of total protein at 2 µg and 0.25 µg per reaction, respectively, and incubated at 4 °C on rotation overnight. Subsequently, Dynabeads™ Protein G magnetic beads (Thermo Fisher Scientific) were added to the lysate/antibody mixture and incubated at RT for 30 min on rotation. Immunoprecipitated proteins were washed 3X with wash buffer [150 mM NaCl, 50 mM Tris-HCl (pH 7.5), 5 % Glycerol] and eluted by an on-bead tryptic digest using elution buffer [2 M Urea (prepared with 50 mM NH_4_HCO_3_ solution), 50 mM Tris-HCl (pH 7.5), 1 mM DTT, 5 µg/ml Trypsin] at 37 °C for 1 h with shaking at 600 rpm. Samples were then alkylated overnight with 5 mM IAA at 37 °C and shaking at 600 rpm. Digested and alkylated peptides were acidified with 1 % TFA, desalted and purified on C18 stage tips for analysis by mass spectrometry.

### Analysis of p62 proxitome by proximity biotinylation-based quantitative proteomics

APEX2-mediated biotinylation of the p62 proxitome in cells was performed by modifying the method described by 39. Briefly, cells were supplemented with 500 mM biotin-phenol (Iris Biotech) for 30 min at 37 °C followed by addition of 1 mM H_2_O_2_ for 1 min at RT. Biotinylation reaction was stopped by washing the cells three times with quencher solution (1 mM sodium azide, 10 mM sodium ascorbate and 5 mM Trolox in DPBS) followed by three washes with PBS. Cells treated with H_2_O_2_ only were included as controls for the biotinylation reaction. Cells were then harvested by scrapping and lysed by dounce-homogenization and sonication in RIPA buffer containing quenching components [50 mMTris, 150 mM NaCl, 0.1 % SDS, 0.5 % sodium deoxycholate, 1 % Triton X-100, 1x protease inhibitors (Merck), 1x PhosStop (Merck), 1 mM sodium azide, 10 mM sodium ascorbate and 1 mM Trolox]. Cleared lysates were obtained by centrifugation at 10000 g for 10 min at 4 °C and incubated overnight on pre-equilibrated Streptavidin-Agarose beads (Sigma). The next day, samples were washed three times with RIPA buffer containing quenching components, followed by three washes in 3 M Urea buffer (prepared with 50 mM NH_4_HCO_3_ solution). Samples were then incubated with 5 mM TCEP for 30 min at 55 °C and alkylated with 10 mM IAA for 20 min at RT. Alkylation was quenched by the addition of 20 mM DTT and samples were washed twice with 2 M Urea buffer (prepared with 50 mM NH4HCO_3_ solution) and digested overnight at 37 °C using 1 µg of trypsin per 20 µl beads. Digested peptides from the supernatants were collected and pooled together with supernatants from two times washes of the beads using 2M Urea buffer. The pooled samples were then acidified with 1 % TFA and concentrated by vacuum centrifugation. Finally, the digested peptides were desalted and purified on C18 stage tips for analysis by mass spectrometry.

### Analysis of p62 cargo candidates engulfed in autophagosomes by proximity biotinylation-based quantitative proteomics

Autophagosomal content profiling for p62 cargo candidates was performed as described by 39. Briefly, APEX2-mediated biotin labeled cells were incubated on an overhead rotator for 20 min at 4 °C in homogenization buffer I (10 mM KCl, 1.5mM MgCl_2_, 10 mM HEPES-KOH and 1mM DTT pH 7.5). Cells were then lysed by dounce homogenization and mixed at a ratio of 1:5 with homogenization buffer II (375 mM KCl, 22.5 mM MgCl_2_, 220 mM HEPES-KOH and 0.5 mM DTT pH 7.5). Cleared lysates obtained by centrifugation at 600 g for 10 min were treated with 100 mg/ml Proteinase K (PK) and 1 mM CaCl_2_ for 1 h at 37 °C, before inhibition of PK activity by adding 10 mM PMSF to the samples. To account for Proteinase K resistant proteins, samples additionally treated with 0.1 % RAPIGest (RP) were added as control. Samples were centrifuged at 17000 g for 15 min at 4 °C. The supernatant was removed, and pellets were resuspended in RIPA buffer containing quenching components, followed by brief sonication and centrifugation at 10000 g for 10 min h at 4 °C. Supernatants were then incubated on pre-equilibrated Streptavidin-Agarose (Sigma) overnight at 4 °C. Samples were then washed three times in RIPA buffer containing quenching components and three times in 3 M Urea buffer (prepared with 50 mM NH_4_HCO_3_ solution). Subsequently, samples were incubated with 5 mM TCEP for 30 min at 55 °C and alkylated with 10 mM IAA for 20 min at RT. Alkylation was quenched by the addition of 20 mM DTT and samples were washed twice with 2 M Urea buffer (prepared with 50 mM NH_4_HCO_3_ solution) and digested overnight at 37 °C using 1 µg of trypsin per 20 µl beads. Digested peptides from the supernatants were collected and pooled together with supernatants from two times washes of the beads using 2M Urea buffer. The pooled samples were then acidified with 1 % TFA and concentrated by vacuum centrifugation. Finally, the digested peptides were desalted and purified on C18 stage tips for analysis by mass spectrometry. To validate the PK protection assay by Western blot, the cleared lysates obtained by centrifugation at 600 g for 10 min were treated with 30 mg/ml PK and 1 mM CaCl_2_ and incubated for 30 min at 37 °C. Control reactions consisting of no digestion, digestions with PK and 0.1 % RP or treatment with 0.1 % RP only were included. Samples were then boiled in Laemmli buffer and subjected to SDS-PAGE.

### Analysis of total proteome by tandem mass tag (TMT)-based quantitative proteomics

Preparation of cell lysates and labelling of peptides with TMT isobaric mass tags was carried out using the commercially available kit and reagents (Thermo Fischer Scientific, #90110) according to the manufacturer’s protocol. Briefly, cell lysates were obtained by suspension of cells in 10 % SDS buffer (prepared with 100 mM triethyl ammonium bicarbonate) followed by sonication. Samples were centrifuged at 16,000 g for 10 minutes at 4 °C and protein concentration determined from the supernatant with a BCA assay (Thermo Fisher Scientific). Approximately 50 µg of protein was reduced with 200 mM TCEP for 1 h at 55 °C and alkylated with 10 mM IAA for 30 min in the dark at RT. Proteins were then acetone precipitated overnight and digested with 1.5 µg proteomics-grade trypsin (Sigma-Aldrich) at 37 °C overnight. Subsequently, 25 µg of digested peptides were labelled with TMT10plex isobaric tags (Thermo Fisher Scientific) for 1 h at RT followed by quenching with 5 % hydroxylamine for15 min at RT. TMT labeled samples were adjusted to equal amounts through a LC-MS test run (with 2 µl of each sample pooled together, desalted, and purified on C18 membranes), before fractionation of the total sample. Individually labelled samples were then pooled, fractionated into 8 fractions using the High pH Reversed-Phase Peptide Fractionation Kit (Thermo Fisher Scientific) according to the manufacturer’s instructions protocol and dried by vacuum centrifugation.

### Liquid chromatography-tandem mass spectrometry

Peptides were resuspended in 9:1 mixture of buffer A (0.1 % formic acid) and buffer A* (2 % acetonitrile, 0.1 % trifluoro acetic acid). Half of the sample was separated on 50 cm columns packed in house with ReproSil-Pur C18-AQ 1.9 µm resin (Dr. Maisch GmbH). Using an EASY-nLC 1200 ultra-high-pressure system connected in-line to a Q-Exactive HF Mass Spectrometer (Thermo Fisher Scientific), liquid chromatography was conducted. Peptides were introduced in buffer A (0.1 % formic acid) and eluted with a non-linear gradient of 5–60 % buffer B (0.1 % formic acid, 80 % acetonitrile) at a rate of 300 nl/min over 90 min. Column temperature was maintained at 60 °C. Data acquisition alternated between a full scan (60 K resolution, maximum injection time of 20 ms, AGC target of 3e6) and 15 data-dependent MS/MS scans (15 K resolution, maximum injection time of 80 ms, AGC target of 1.6e3). The isolation window was 1.4 m/z, and normalized collision energy was 28. Dynamic exclusion was set to 30 seconds. TMT-labelled peptides were separated as described above except that a 180 min gradient was used for elution. MS data was acquired in data-dependent mode using the following settings. Full MS scans (120 K resolution, maximum injection time of 80 ms, AGC target of 3e6) were alternated with a set of maximally 15 data-dependent MS/MS scans (60 K resolution, maximum injection time of 100 ms, AGC target of 2.0e3). The isolation window was 0.7 m/z, and normalized collision energy was 32. Fixed first mass was set to 100 m/z. Dynamic exclusion was set to 30 seconds.

### Mass spectrometry data analysis

Raw mass spectrometry data was processed using using MaxQuant (2.0.1.0). Peak lists were searched against a Uniprot database (UP000005640 9606), alongside 262 common contaminants using the Andromeda search engine. A 1 % false discovery rate was applied for both peptides (minimum length: 7 amino acids) and proteins. The “Match between runs” (MBR) function was activated, with a maximum matching time window of 0.7 min and an alignment time window of 20 min. Relative protein quantities were calculated using the MaxLFQ algorithm, requiring a minimum ratio count of two. The calculation of iBAQ intensities was enabled.

### Computational analysis of mass spectrometry data

The protein groups identified in each mass spectrometry data set were processed and analyzed in parallel using the R programming language (version 4.3.3; R Core Team 2024). First, the MaxQuant output was filtered to remove contaminants, reverse hits, proteins identified by site only as well as proteins that were identified in less than N-1 replicates of every condition, where N is the number of replicates (*N* ≥ 3). The label-free quantification (LFQ) or corrected reporter ion intensities of the robustly identified proteins were log2-transformed subsequently. For the global proteome data set only, the protein intensities were normalized between samples using fast cyclic loess normalization as implemented in the R package *limma* (*69*) (version 3.58.1). Lastly, missing values were imputed by sampling N values from a normal distribution (seed = 12345) and using them as replacements only when there are no valid values in a group of replicates of a condition or if there is only one valid value which is below the first tertile of intensities in the data set. Different for each data set, the mean of this normal distribution corresponds to the 1 % percentile of LFQ intensities, and its standard deviation is determined as the median of protein intensity sample standard deviations calculated within and then averaged over each group of replicates.

Principal component analysis was carried out for each data set using the package *pcaMethods* (*70*) (version 1.94.0) on the processed and standardized protein intensities.

Protein groups were statistically analyzed using pairwise, two-sided Student’s *t*-test s on the processed protein intensities between the replicates of respective control and test conditions. Only proteins with valid values in the test conditions were considered for analysis. Log2 fold changes were derived as the tested difference of means and the resulting *P*-values were adjusted for the false discovery rate (FDR) using the Benjamini-Hochberg procedure.

Using the results of our co-IP mass spectrometry data analysis we define robust interactors of p62 per treatment background as those proteins, whose intensities differ significantly (FDR *<* 0.05) between the p62 pulldown in a specific cell line and all other groups of IgG pulldown controls individually. Furthermore, to be able to determine the exclusivity of p62 interactors between each karyotype, we tested whether a protein could be either classified as an interactor using the conservative definition above or be excluded as an interactor by not meeting the more inclusive condition of a significant difference in intensities between the p62 pulldown in a cell line and only the matching IgG control with a *P*-value *<* 0.05. Subsequently, we only considered candidate interactors that fulfill this requirement for down-stream analysis.

The results from the proximity biotinylation experiments were adjusted to account for alterations in overall protein levels across whole-cell extracts. We used the comprehensive proteomic dataset for HCT116 5/4 cells (*8*), while data obtained in this work were used for HCT116 13/3. Subsequently, the Student’s *t*-test s were re-evaluated using the global log2 fold changes as the true difference in revised means of the new null hypothesis. The proxitome fold changes were corrected by subtracting the global log2 fold changes. The analyses reflect whether the difference in protein intensities relative to the wild type is significantly higher within the p62 proximity than in the global abundance.

### Over-representation analysis

Hypergeometric tests were performed to assess the statistical significance of the overlap between the sets of differentially abundant proteins and the different molecular signature sets given all robustly identified proteins as a background. FDR-adjustment of *P*-values was done according to Benjamini-Hochberg within signature set categories. When indicated, the redundancy be-tween related sets was reduced by applying the affinity propagation algorithm with negative log-transformed *P*-values for scoring to cluster similar sets and extract representatives as implemented in the *WebGestaltR* package (*71*) (version 0.4.6).

### Deriving an autophagy related gene set

A set of autophagy related genes was derived by filtering all gene sets in the Molecular Signature database (*72*) for sets with names including “AUTOPHA” or “AUTOLYSOSOME”. This way, we identified and combined 40 gene sets from sources including Reactome (*73*), KEGG (*74*), Gene Ontology (*40, 41*) and WikiPathways (*75*) gene sets to obtain a set of 257 autophagy related proteins (see Supplementary Table 3).

### Hydrophobicity scores

The function scaledHydropathyLocal implemented in R package *idpr* (*76*) (version 1.12.0) was applied to calculate the scaled hydropathy of all amino acid sequences in the UniProt database (UP000005640 9606) using a sliding window of 21 amino acids. Hydrophobicity of a protein group identified in the mass spectrometry proteomics experiments were then derived as the average maximum scaled hydropathy of all proteins in a group.

### Analysis of external cancer cell line multi-omics data

Cancer cell line data from the DepMap database was filtered to only include cancer cell lines that did not undergo whole genome doubling (WGD) as previously inferred (*36*). This was done to exclude any confounding effects of WGD positive cancer cell lines and to make the selection of cancer cell lines more comparable to our WGD negative HCT116. This left 127 cancer cell lines with both aneuploidy scores and transcriptome, proteome and gene dependency data. Subsequently, cell line gene expression and dependency values were correlated with cell line aneuploidy scores using Spearman’s rank correlation coefficient as implemented in the base R-package.

### Visualization of omics data

The results of the computational analysis of mass spectrometry data and the analysis of external cancer cell line multi-omics data were visualized using the *ggplot2* (*77*) (version 3.5.0), *ggpubr* (version 0.6.0), ggh4x (version 0.2.8) and *cowplot* (version 1.1.3) R packages.

### Cell lysis, SDS-PAGE and Western blot

Whole-cell lysates were prepared by solubilizing cell pellets in RIPA buffer supplemented with Protease and Phosphatase inhibitors (Merck) and vortexing every 5 min for 30 min at 4 °C. Samples were centrifuged at 2000 g for 10 mins to obtain the protein supernatant from which protein concentration was determined by the Bradford assay. To prepare samples for SDS-PAGE, lysates were diluted to 1 µg/µl with Laemmli buffer and boiled for 5 min at 95 °C. 10 µg of total protein was run on SDS-PAGE gels using either the Precision Plus Protein All Blue Standard (Bio-Rad) or Color Prestained Protein Standard (New England Biolabs) as marker. Proteins were transferred onto nitrocellulose membranes (AmershamProtran Premium0.45 NC, GE Healthcare Life Sciences, Sunnyvale, USA) by wet transfer at 100 V for 1 h and blocked in 3 % BSA solution for 1 h at RT. Primary and secondary antibody incubations were carried out overnight at 4 °C and 1 h at RT, respectively. Protein signals were visualized by chemiluminescence and imaged on an Azure c500 system (Azure Biosystems, Dublin, USA). Quantification of protein bands was done in ImageJ. A list of antibodies used can be found in Supplementary Table 5.

### Immunofluorescence

Cells were grown on poly-L-lysine coated coverslips prior to the indicated treatments. Thereafter, cells were washed three times using PBS and fixed using a 2 % PFA/ 2 % sucrose solution for 10 min at RT. Fixed cells were then washed three times using PBS, permeabilized using 0.1 % TX-100 and blocked with 3 % BSA for 1 h at RT. Primary and secondary antibodies were incubated overnight at 4 °C, followed by 1 h at RT in the dark, respectively. Coverslips were mounted on microscopy slides using VECTASHIELD® Hardset Antifade Mounting Medium containing DAPI (Vector Laboratories Wertheim, Germany). Slides were imaged using a semi-automated Zeiss AxioObserver Z1 (Oberkochen, Germany) equipped with an ASI MS-2000 stage (Applied Scientific Instrumentation, Eugene, USA), a CSU-X1 spinning disk confocal head (Yokogawa), a Laser Stack with selectable lasers (Intelligent Imaging Innovations, Denver, USA) and the Cool-Snap HQ camera (Roper Scientific). 40x air or 63x oil objectives were used under the control of the Slidebook6 software (Intelligent Imaging Innovations, Denver, CO).

### Microscopy image analyses

p62-positive foci number and size were analyzed using Z-projections of p62-immunostained cells. Cell boundaries (i.e., regions of interest or ROI) were set manually by applying adjustments in brightness and contrast to the images in p62 channel. A threshold was set to reduce signal-to-noise ratio by using background signal from the WT cells as a reference. Additionally, a second threshold was set using the “Analyze particles” function in ImageJ to exclude structures less than 0.01 µm in size as p6-positive foci. Foci were then evaluated using an in-house automated pipeline developed based on ImageJ macros language. To account for differences in cell sizes, foci counts were normalized to the cells surface area estimated from the ROIs.

Colocalization of p62 and TOMM20 was estimated using the Ez-Colocalization plugin in ImageJ (*78*). A maximal intensity projection on the Z-axis was performed and this single in-focus field projection was used for the colocalization analysis. In the EzColocalization plugin, an automated threshold was set for the TOMM20 signal, whilst threshold of the p62 channel was set using background signal from the p62 channel in WT cells. Individual cells were outlined and saved to the ROI manager tool in ImageJ and analyzed with the “ROI” cell identification input. The output data was selected as Pearson correlation coefficient. This value ranges between -1 and 1. A perfect colocalization event is signified by a perfect linear correlation of fluorescence intensities of two images and assumes a Pearson correlation coefficient value of 1. A negative linear correlation results in a value of -1, and no correlation results in a value of 0 (*79*).

To evaluate mitochondrial network, the Mitochondria Analyzer plugin (*80*) was used in ImageJ. The threshold algorithm of this plugin requires the empirical determination of block size and C-value parameters by a user, which prevents the merging of close but physically separate individual mitochondria as well as fake splitting of continuous mitochondria. An optimal threshold for the TOMM20 channel was determined to be C value of 1 µm and block size of 1.45 µm, which gave the most accurate separation of the mitochondrial network in all cell lines used. Individual non-overlapping TOMM20 immunostained cells were then cropped out following the excision of mitochondria from adjacent cells where necessary. Mitochondria morphology analysis was performed on “Per-cell basis” and mean values of the following parameters were obtained: Branches, Total Branch Length, Branch Junctions, Branch Points, Mitochondrial Volume, and Mitochondrial Surface Area. Classification of network as extended, intermediate and perinuclear was determined using the number of branch junctions.

### Flow cytometry

Cultured cells were stained simultaneously with MitoTracker Green and Deep Red dyes according to the manufacturer’s instructions to analyze mitochondria mass and membrane potential, respectively, and incubated for 20 min at 37 °C and 5 % CO_2_. Unstained cells were included for background control. After incubation, cells were washed five times with PBS before harvesting by trypsinization. Cells were then resuspended in PBS and analyzed on an Attune NxT acoustic focusing cytometer (Thermo Fisher). The gating, data analysis, and visualization were performed using the FlowJo software (version 10).

### Mitochondria isolation and import assay

Isolation of mitochondria and protein import assay were performed as previously described (*55*). Briefly, confluent cells were harvested and washed with ice-cold 1X PBS. Cells were then resuspended in homogenization buffer (220 mM mannitol, 70 mM sucrose, 5 mM HEPES-KOH pH 7.4, 1 mM EGTA-KOH pH 7.4) containing 1× Complete Protease Inhibitor Cocktail (Roche). and homogenized using a PTFE-based homogenizer (VWR). Homogenate was clarified by centrifugation at 600 g for 5 min at 4°C to remove cell debris and nuclei. The supernatant was further centrifuged at 8,000 g for 10 min at 4°C to obtain the crude mitochondrial fraction, which was washed once in homogenization buffer before the import assay. Subsequently, import assay was performed with 20 µg of mitochondria per reaction. Radiolabeled precursor proteins were synthesized using the SP6 promoter TNT Quick Coupled Transcription/Translation System (Promega) containing 20 µCi [^35^S]-methionine at 30 °C for 1 h. The import reaction was started by incubating precursor protein with crude mitochondria at 30 °C in the presence or absence of 1 mM CCCP. Precursor protein import was then stopped after 1, 2, 5, 10 or 15 min by placing mitochondria on ice. Proteinase K treatment was performed to degrade unimported substrates. The mitochondria were then washed in homogenization buffer containing 1 mM PMSF, resuspended in Laemmli buffer and analyzed by SDS–PAGE followed by autoradiography.

### Mitochondrial oxygen consumption assay

100 µg of isolated intact mitochondria was resuspended in a respiration buffer containing 0.6 M sorbitol, 20 mM HEPES/KOH (pH 7.4) and 1 mM MgCl_2_ to determine respiration capacity. The mitochondrial oxygen consumption rate was measured using a Clark-type oxygen electrode connected to a computer-operated Oxygraph control unit (Hansatech Instruments, Germany) at 25 °C with continuous stirring. The upper (100 %) and lower (0 %) limit of dissolved oxygen was calibrated using distilled water and sodium dithionite, respectively. Mitochondria were added to the chamber of the unit and respiration was initiated by the addition of 7 mM NADH. Data was recorded at intervals of 1 s (Oxygraph Plus software, Hansatech Instruments). In a control reaction, isolated mitochondria were treated with 10 mM CCCP for 15 mins to collapse the membrane potential before oxygen consumption measurements. Oxygen consumption (in µmol/µl) was normalized to the mitochondrial protein amount determined by Bradford assay.

### Quantification of mitochondrial DNA copy number by qPCR

Total DNA was extracted from cultured cells using the QIAamp DNA Mini Kit according to the manufacturer’s protocol. Mitochondrial DNA was quantified from 10 ng of total DNA using SYBRGreen Mastermix and primer sets that amplify the non-coding D-Loop region and coding sequences of at least one respiratory chain complex subunit (Supplementary Table 6). The nuclear encoded ß_2_ microglob-ulin gene was used for normalization. Samples were assayed on the C1000 Touch Thermal Cycler (Bio-Rad) using the following run conditions: initial denaturation for 5 min at 95 °C followed by 45 cycles of 5 s at 95 °C, 15 s at 55 °C, 15 s at 72 °C and 1 s at 78 °C.

### Statistical analyses

Outside the analysis of omics data, Graph-Pad Prism 9 was used for statistical analysis. For each data set, the Kolmogorov-Smirnov-test was applied to determine whether variables follow a normal distribution. If yes, a two-tailed unpaired *t*-test was used to compare the means of two groups. For multiple comparisons, ordinary one-way ANOVA was used followed by Sidak’s post hoc test. For non-normally distributed data, one-way ANOVA on ranks (Kruskal-Wallis H Test) was performed followed by Dunn’s post hoc test.

## Data availability

The mass spectrometry proteomics data (see also Supplementary Tables 1, 2, 3) have been deposited to the ProteomeXchange Consortium via the PRIDE partner repository with the dataset identifiers PXD052174, PXD052623, PXD052624 and PXD052637 and are available upon reasonable request. Multi-omics data of CCLE cancer cell lines, including aneuploidy scores (*34*), RNAseq transcriptomics (*36*), mass spectrometry proteomics (*37*) and gene dependency (*35*) data were downloaded through the DepMap data portal (*32*) (version 22Q4). Gene Ontology Cellular Compartments (GOCC) and other gene sets were downloaded from the Molecular Signature database (*72*) (version 2022.1). Information on macromolecular complexes were downloaded from the CORUM database (*81*). Scripts used to analyze the data and generate the figures are available upon request.

## Extended Data Figures

**Extended Data Fig. 1.**
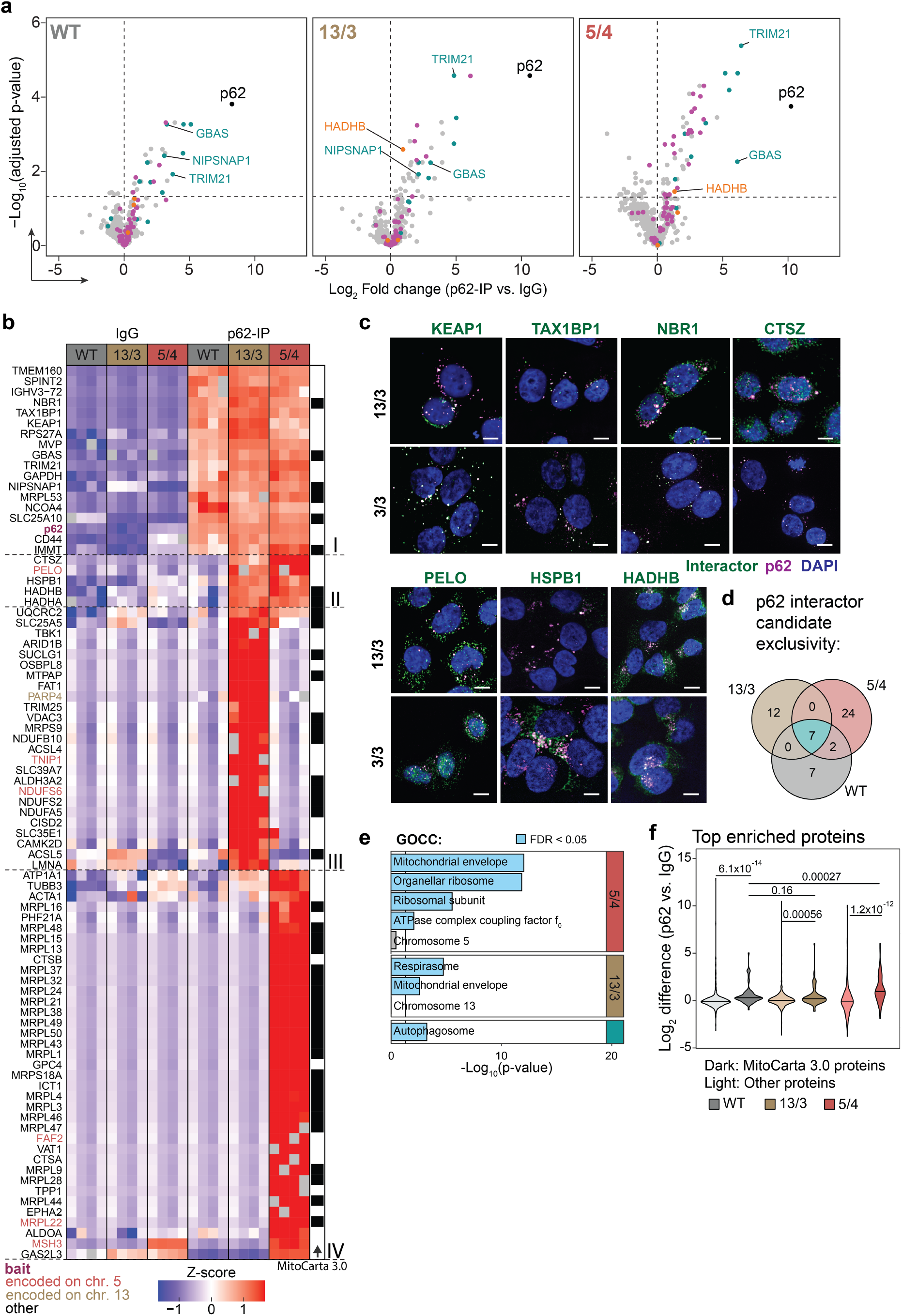
Analyses of p62 interactome by IP-MS. **a**, Volcano plots showing log2-transformed fold change of enriched p62 interactors in DMSO (vehicle) treated WT, 13/3 and 5/4 cells highlighting mitochondrial (magenta), karyotype-independent (cyan) and polysomic-specific (orange) interactors. **b**, Heatmap of hierarchical clustering of enriched p62 interactors from (**Fig. 2d**) according to their Z-scores. Black boxes represent mitochondrial proteins as defined by MitoCarta 3.0, clusters I – IV present proteins with similar pattern of enrichment. Missing values are shown in grey. **c**, Representative confocal images of colocalization of p62 with selected interactors in 13/3 and 3/3 cells. Scale bar 10 µm. **d**, Venn diagram showing distribution of enriched p62 interactors between karyotypes when compared to IgG control from all groups in vehicle (DMSO)-treated cells. **e**, Gene ontology (GO) over-representation analysis of enriched p62 interactors in (**a**). Blue bars represent negative log-transformed *P*-values for GO terms with FDR *<* 0.05. Representatives from clusters of related GO terms are shown (see Methods). **f**, Comparison of fold differences in enrichment of mitochondrial proteins, as defined by the MitoCarta 3.0 inventory, to all other proteins in the measured p62 interactome in (**a**). *P*-values are derived from two-sided Wilcoxon’s ranks sum tests.

**Extended Data Fig. 2.**
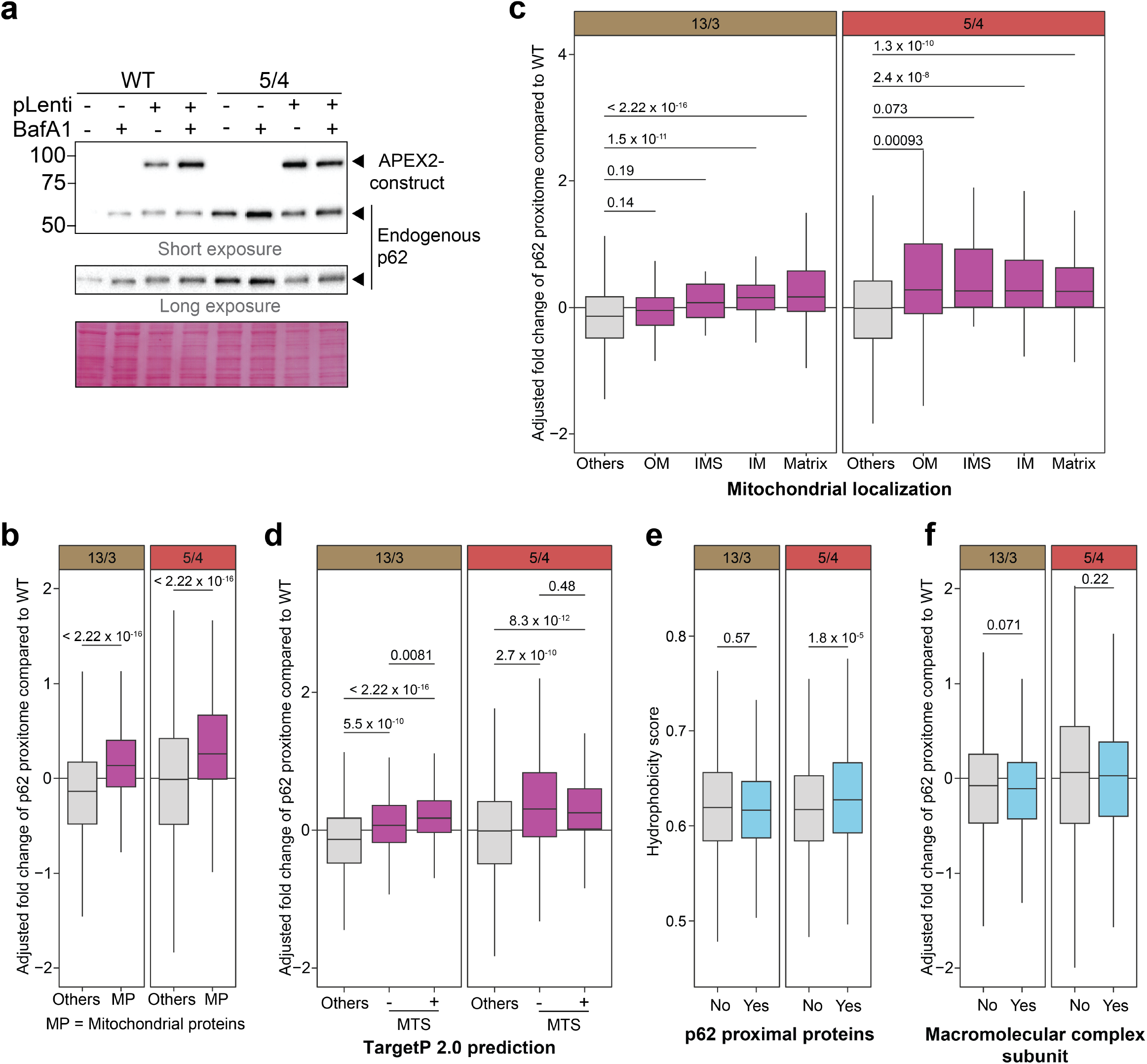
Characterization of p62 proximal proteins in polysomic cells. **a**, Immunoblot showing stable expression of the APEX2-p62 construct in WT and 5/4 cells in the absence and presence of BafA1. The expression of endogenous p62 was also monitored. Ponceau staining is used as loading control. **b**, Boxplots indicating adjusted fold changes in abundance of mitochondrial proteins (magenta) compared to all other measured proteins. **c**, Fold change distribution of p62-proximal mitochondrial proteins according to subcellular localization. **d**, distribution of the enriched p62-proximal mitochondrial proteins based on whether they possess canonical mitochondrial targeting sequences (MTS) according to TargetP 2.0 prediction. **e**, Hydrophobicity score, and **f**, Identity as subunit of macromolecular complexes (CORUM) of the enriched p62-proximal proteins. *P*-values are derived from two-sided Wilcoxon’s ranks sum tests.

**Extended Data Fig. 3.**
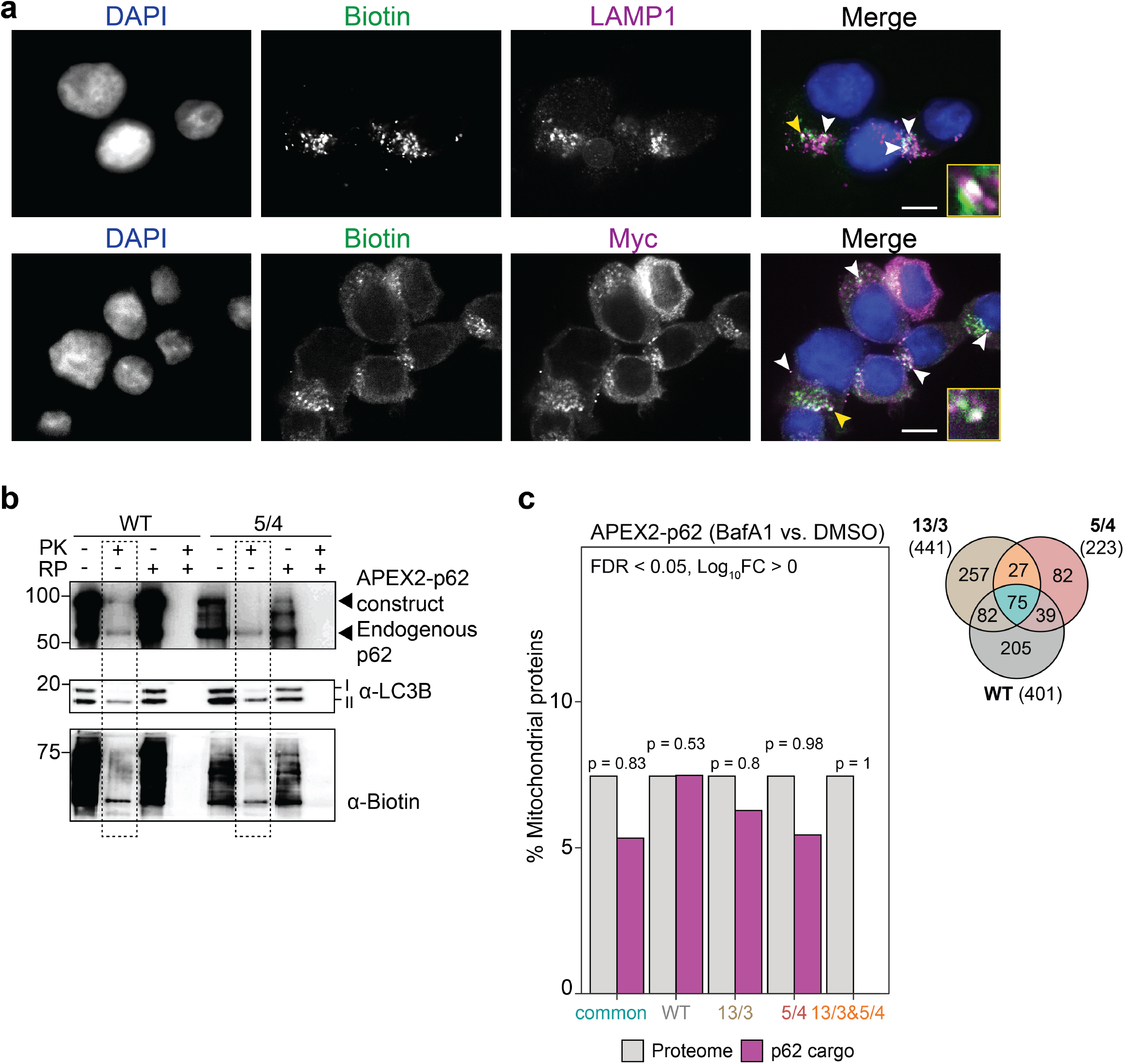
Analyses of autophagosome luminal p62 proximal cargo. **a**, Representative confocal images indicating colocalization of biotin with APEX2-p62 and the autolysosomal marker LAMP1. Scale bar, 10 µm. Colocalization events are indicated by white and yellow arrowheads. Insets represent colocalizations indicated by yellow arrowheads. **b**, Immunoblot of the protease protection assay. Cell homogenates were incubated with Proteinase K (PK), RAPIGest (RP), or both and blotted for p62, LC3B and biotin. Dotted box show condition analyzed by mass spectrometry. The presence of only LC3B-II in this condition confirms successful enrichment of autophagosomes. **c**, Percentage of mitochondrial proteins in the measured lumen proteome (*n* = 1610) and the different sets of p62 cargo candidates as indicated in the Venn diagram, which shows the distribution of the number of autophagosomal p62 cargo candidates between karyotypes. *P*-values represent results of hypergeometric test, evaluating the differences in representation of mitochondrial proteins.

**Extended Data Fig. 4.**
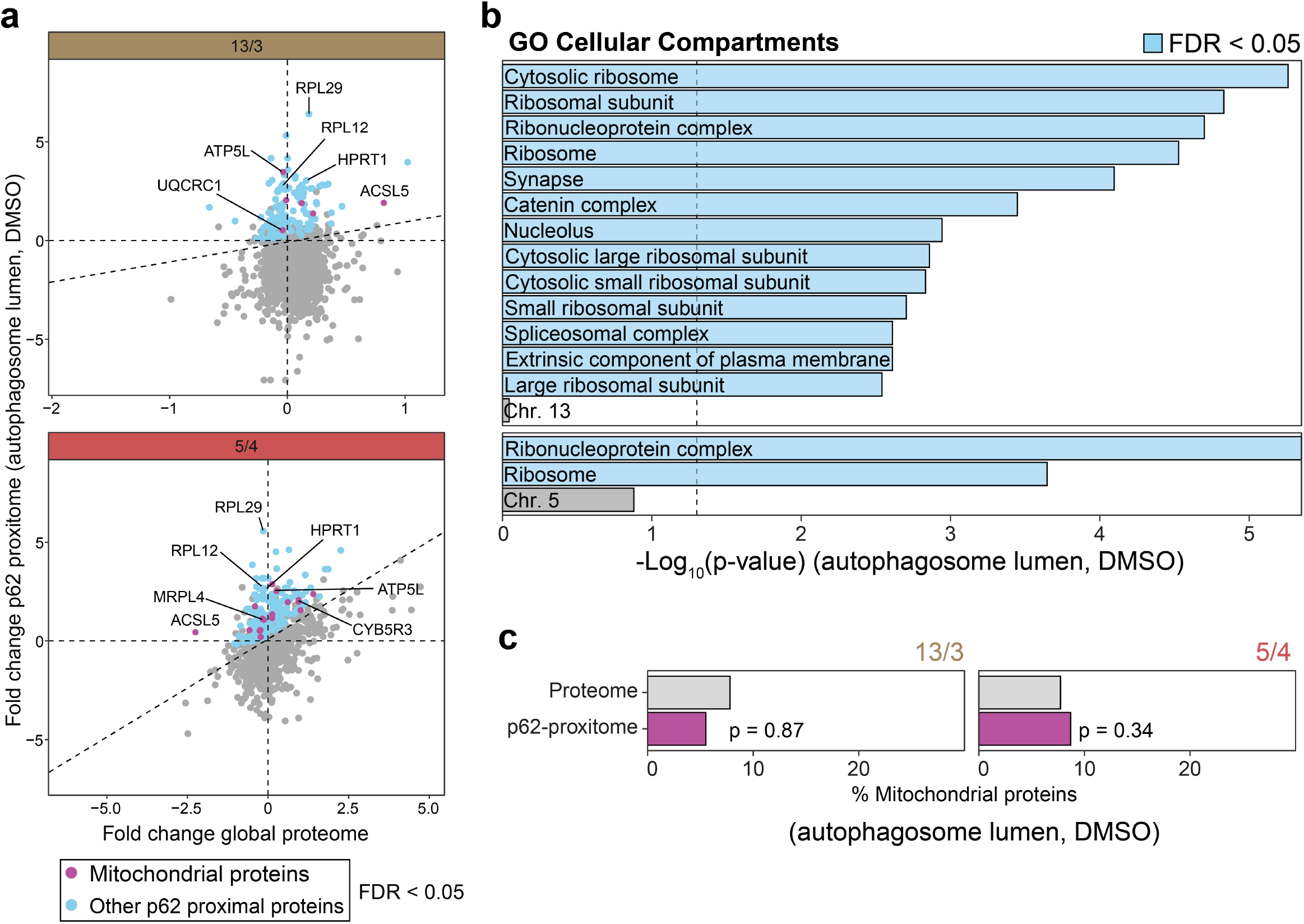
Autophagosome content profiling reveals ribosome and ribonucleoproteins as predominant p62 proximal proteins in autophagosome lumen of polysomic cells. **a**, Scatter plot of the fold changes in global protein abundance in untreated 13/3 and 5/4 polysomic cell lines relative to the parental cell line (WT) and the corresponding changes in abundance of the autophagosome lumen p62-proximal proteins. Colored dots represent proteins with significantly higher abundance changes of autophagosome lumen p62-proximal proteins in polysomic cells (FDR *<* 0.05). **b**, Gene ontology (GO) over-representation analysis of the enriched p62-proximal proteins in (**a**). Blue bars represent negative log-transformed *P*-values for GO terms with FDR *<* 0.05. **c**, Percentage of mitochondrial proteins in the measured lumen proteome (*n* = 1610) and the corresponding percentage within the proteins increased in the autophagosome lumen p62 proximal proteome of 13/3 and 5/4 cells. *P*-values represent results of hypergeometric test, evaluating the differences in representation of mitochondrial proteins.

**Extended Data Fig. 5.**
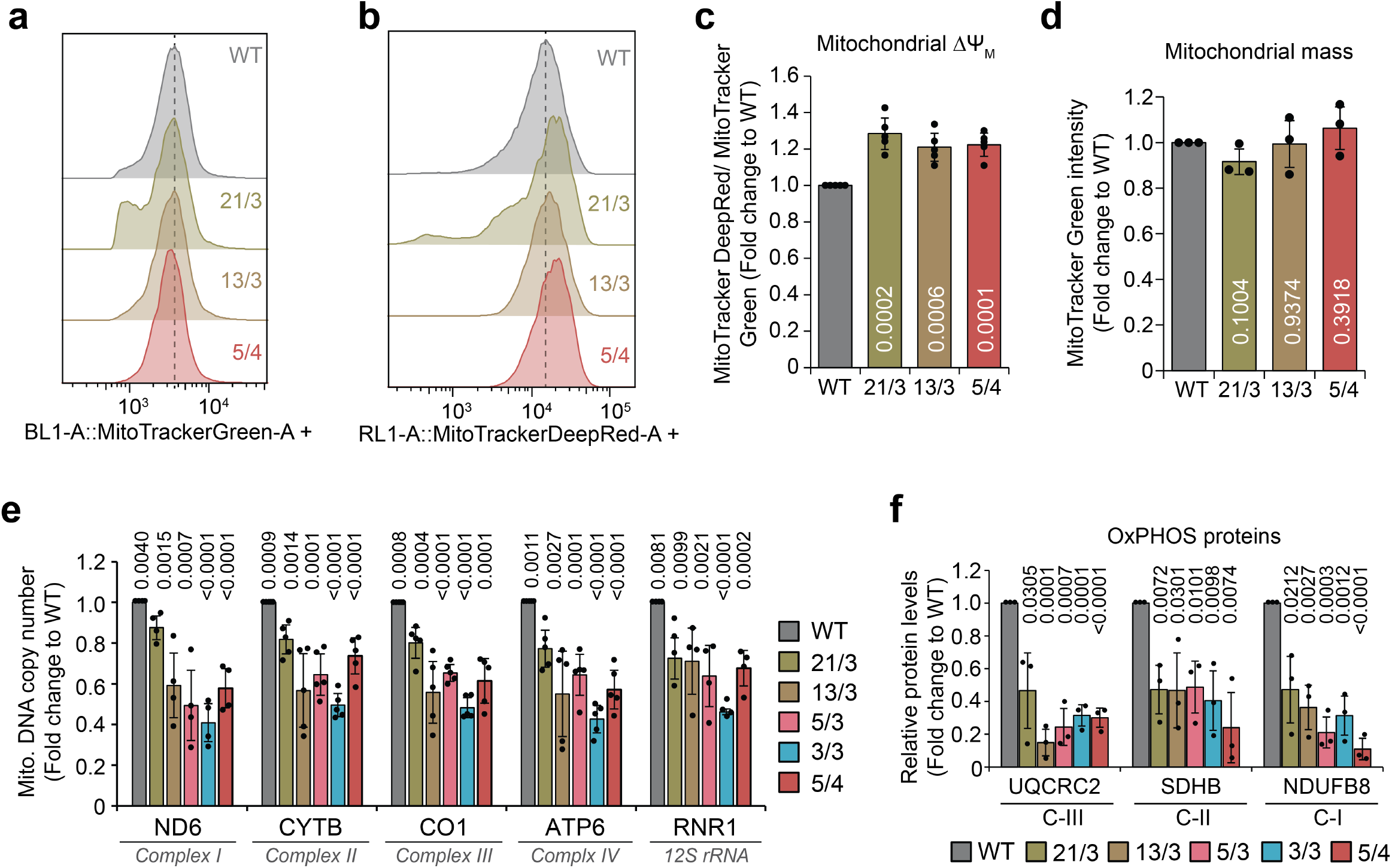
Analyses of mitochondrial phenotypes. **a**, Representative histograms of Mitotracker Green FM and **b**, Mitotracker DeepRed FM fluorescence measured by flow cytometry. **c**, Mitochondrial membrane potential quantified as ratio of Mitotracker DeepRed FM to Mitotracker Green FM intensity. **d**, Mitochondrial mass determined from Mitotracker Green FM intensities. Data is shown as mean ± s.d. fold change to WT from at least *n* = 3 independent experiments, and individual replicates are shown as dot plots. *P*-values represent two-tailed unpaired Student’s *t*-test. **e**, Relative mitochondrial DNA (mt-DNA) copy number determined by qPCR of mitochondrially encoded respiratory chain complex subunits, normalized to the nuclear ß_2_-microglobulin gene. Data is shown as mean ± s.d. fold change to WT from *n* = 5 independent experiments, and individual replicates are shown as dot plots. *P*-values represent two-tailed unpaired Student’s *t*-test. **f**, Quantification of expression levels of selected mitochondrial proteins from (**Fig. 5g**). Data is shown as mean ± s.d. fold change to WT from at least *n* = 3 independent experiments, and individual replicates are shown as dot plots. *P*-values represent two-tailed unpaired Student’s *t*-test.

**Extended Data Fig. 6.**
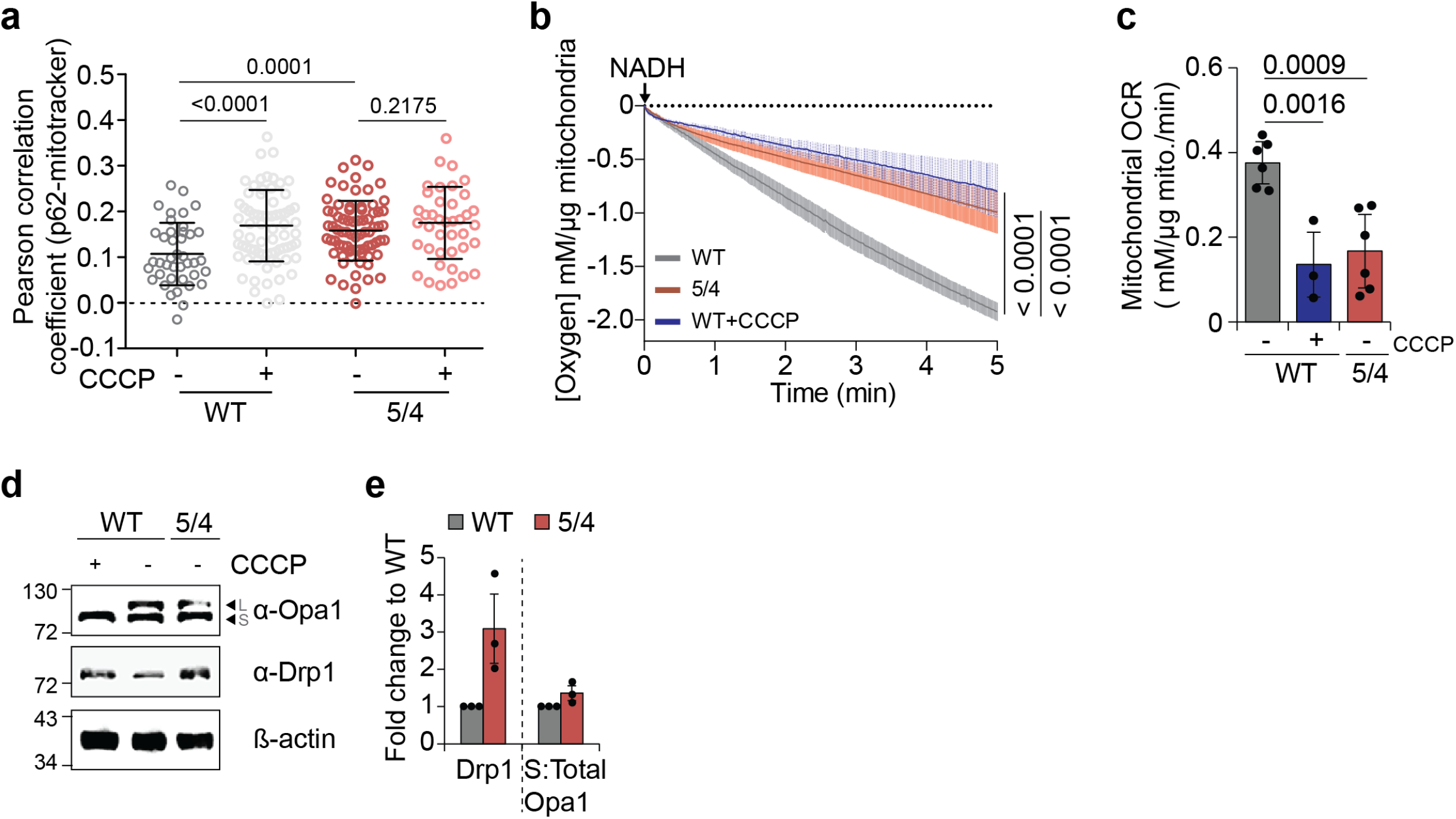
Comparative analyses of mitochondrial functions. **a**, Quantification of the Pearson correlation coefficient for p62-TOMM20 colocalization in WT and 5/4 cells upon DMSO (vehicle) and 10 µM CCCP treatment for 1 h. Individual data of cells from *n* = 3 independent experiments (dot plot) and mean are shown. Error bars represent standard deviation. *P*-values represent one-way ANOVA followed by Sidak’s multiple comparisons test. **b**,NADH-induced oxygen consumption (respiration) of mitochondria isolated from WT and 5/4 cells. In a control reaction, isolated mitochondria from WT cells were treated with 10 mM CCCP for 15 min prior to oxygen consumption measurement. Data is mean ± s.e.m. of at least *n* = 3 independent assays. *P*-value represent one-tailed paired *t*-test. **c**, Oxygen consumption rate (OCR) determined from (**b**). *P*-values represent two-tailed unpaired Student’s *t*-test. **d**, Representative immunoblot and **e**, quantification of protein bands from (**d**). ß-actin is loading control. Data is shown as mean ± s.d. fold change to WT from *n* = 3 independent experiments, and individual replicates are shown as dot plots.

**Extended Data Fig. 7.**
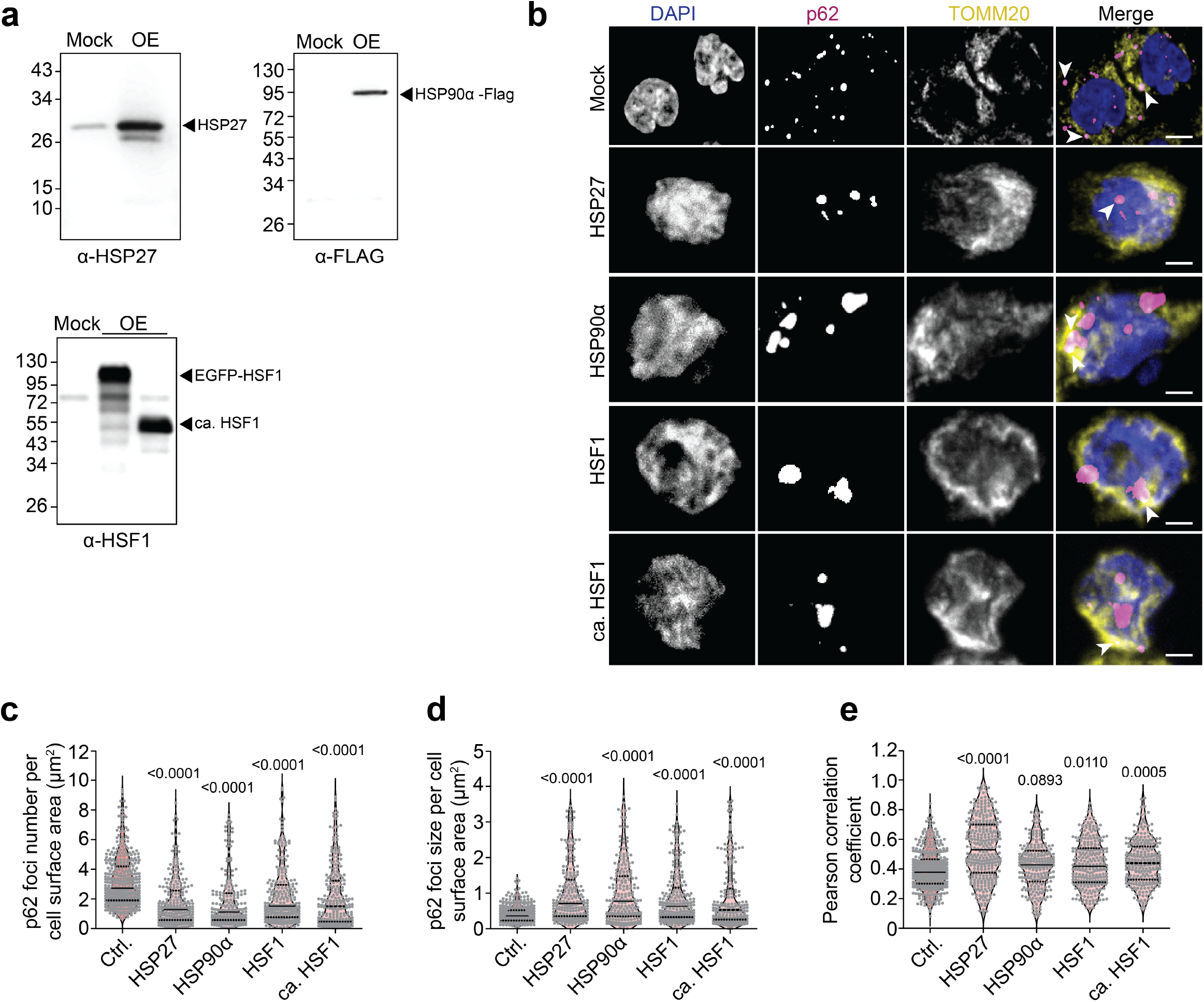
Transient overexpression of heat shock factors in polysomic cells. **a**, Representative immunoblot showing overexpression of HSP27, HSP90α-Flag, EGFP-HSF1 and constitutively active HSF1 in 5/4 cell lines. **b**, Representative confocal images of p62 foci and TOMM20 visualization in the HCT116 5/4 cells overexpressing heat shock proteins. Scale bar 10 µm. **c**, Quantifications of p62 foci number and **d**, p62 foci size per cell surface area in µm^2^ in the analyzed samples. Violin plots indicate mean and distribution of data while dot plots represent individual cells from *n* = 3 independent experiments. *P*-values represent non-parametric ANOVA (Kruskal–Wallis statistic for (**c**) 195.4, p *<* 0.0001; for (**d**) 137.3, p*<* 0.0001) followed by Dunn’s multiple comparisons test. **e**, Pearson correlation coefficient for p62-TOMM20 colocalization in samples from (**b**). Violin plots indicate mean and distribution of data while dot plots represent individual cells from *n* = 3 independent experiments. *P*-values represent one-way ANOVA followed by Sidak’s multiple comparisons test.

**Extended Data Fig. 8.**
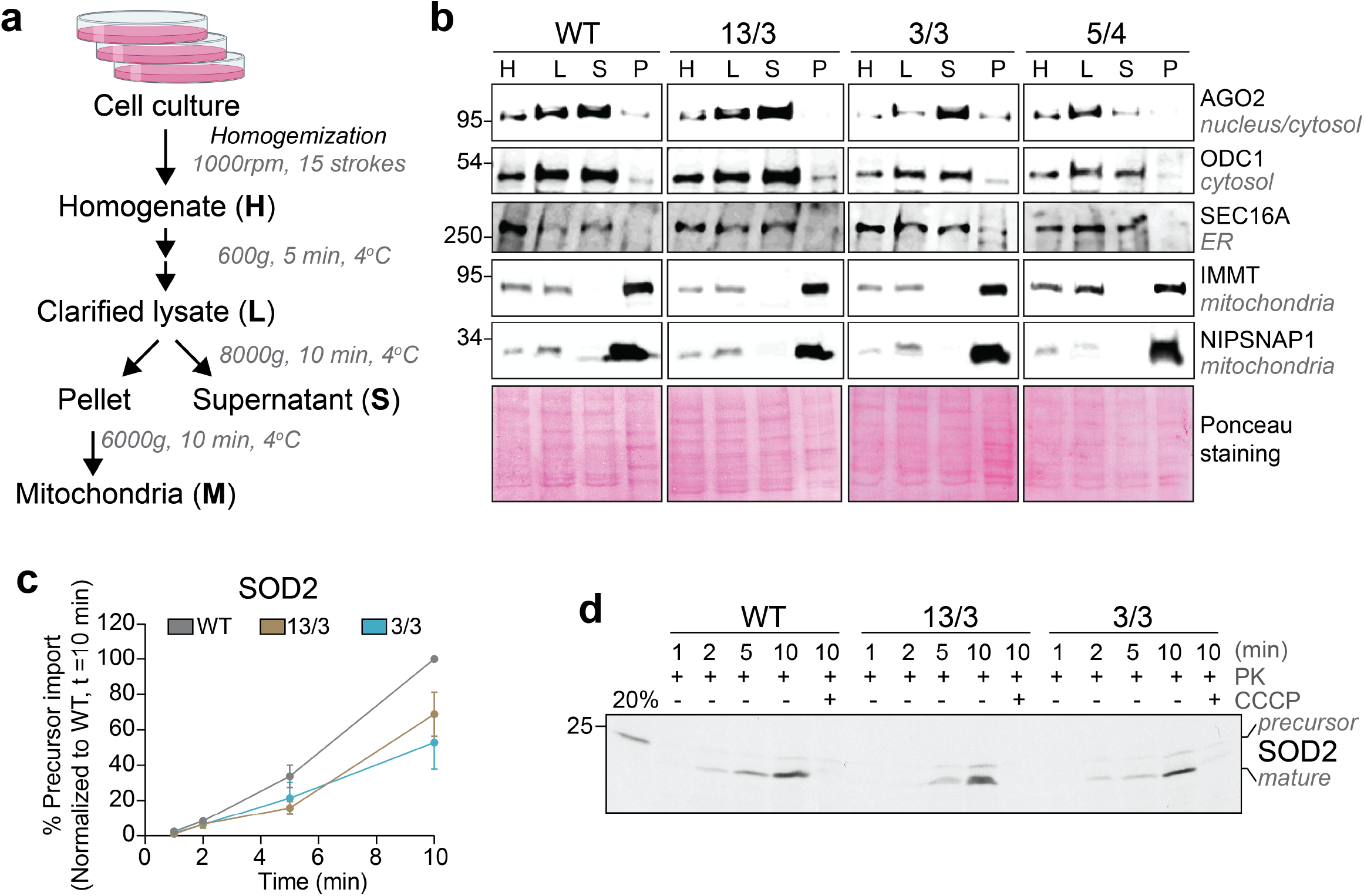
Protein import into isolated mitochondria. **a**, Schematic representation of procedure for mitochondrial isolation from human cells by differential centrifugation. **b**,Representative immunoblots to control for mitochondrial isolation in (**a**) from all the cell lines used. **c**, Quantification and **d**, Representative image of import kinetic of *in-silico* synthesized ^35^S-methonine-labeled human SOD2 into mitochondria isolated from WT, 13/3, and 3/3 cells. The SOD2 precursor is processed upon reaching the matrix (i.e., mature form). CCCP depletes mitochondrial membrane potential thereby preventing import. 20 % of the synthesized substrate (precursor) was loaded for comparison. All samples were resolved by SDS-PAGE and visualized by autoradiography. Data represents mean ± s.e.m. of *n* = 3 independent import assays.

## Supplementary Data Figures

**Supplementary Data Fig. 1.**
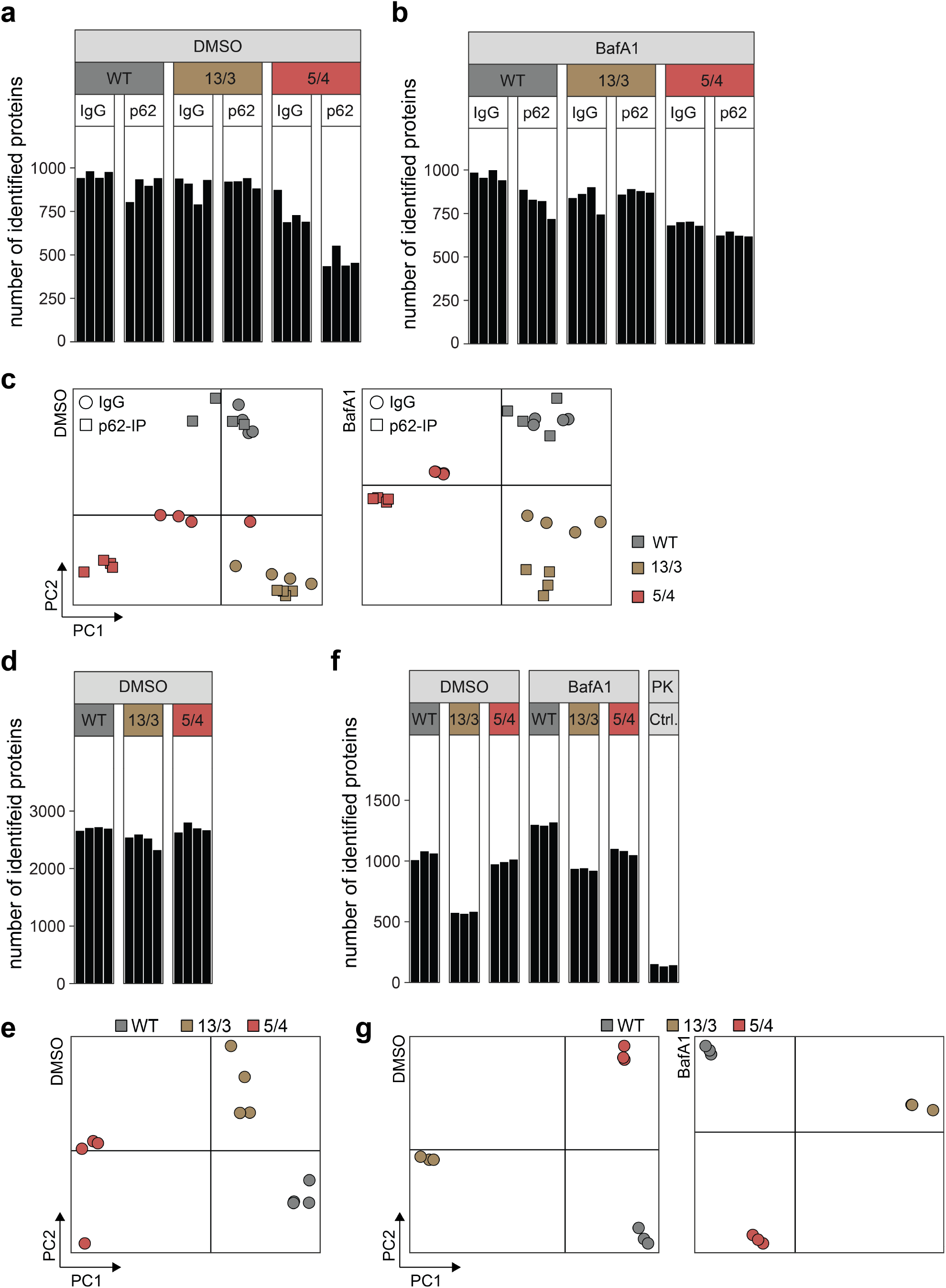
Chromosome gain impairs mitochondrial precursor protein import. Number of proteins from proteomics data of IP-MS in DMSO and **b**, Bafilomycin A1 treated cells. **c**, PCA of proteomics data from (**a-b**). **d**, Number of proteins from cytosolic p62 proximal proteome of DMSO and Bafilomycin A1 treated cells. **e**, PCA of proteomics data from (**d**). **f**, Number of proteins from autophagosome lumen p62-proximal proteome of DMSO and Bafilomycin A1 treated cells. **g**, PCA of proteomics data from (**f**). PK Ctrl. = proteinase K resistant control (clarified lysates treated with both proteinase K and RAPIGest to identify proteins not digested by proteinase K).

**Supplementary Data Fig. 2.**
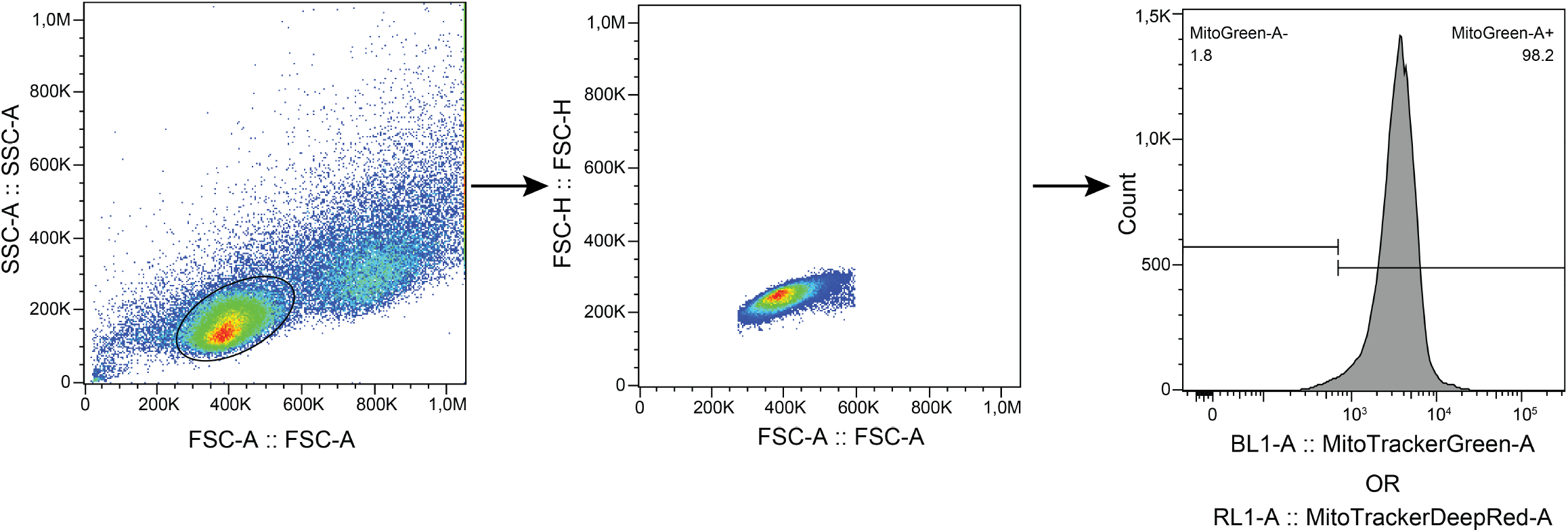
Gating strategy for Mitrotracker stained cells.

