## Supplementary Table 4-6 for "Aneuploidy-induced proteostasis disruption impairs mitochondrial functions and mediates aggregation of mitochondrial precursor proteins through SQSTM1/p62"

**Supplementary Table 4:** List of plasmids

| **Name** | **Purpose** | **Source** |
| --- | --- | --- |
| pHDM-Hgpm2 | Lentivirus packaging | Brass et al. 2008 |
| pHDM-tat1b | Lentivirus packaging | Brass et al. 2008 |
| pHDM-VSV-G | Lentivirus packaging | Brass et al. 2008 |
| pRC-CMV-rev1b | Lentivirus packaging | Brass et al. 2008 |
| pHAGE-myc-APEX2-p62 | Stable transfection | Zellner et al. 2021 |
| pCDNA3.1 HSP90α-Flag | Transient transfection | Kind gift from Len Neckers |
| pCDNA3.1+ ca. HSF1 | Transient transfection | Kind gift from Ulrich Hartl |
| pEGFP-N2 HSF1 | Transient transfection | Kind gift from Ulrich Hartl |
| pCMV5 HSP27 | Transient transfection | Kind gift from Bianca Brundel |
| pEGFP-N3 | Transient transfection | Kind gift from Anne Simonson |
| pEGFP-p62 | Transient transfection | Lamark et al. 2003 |

**Supplementary Table 5:** List of antibodies

| **Product** | **Species** | **Supplier** | **Product number** |
| --- | --- | --- | --- |
| Anti p62 (SQSTM-1) Ick ligand | Mouse | BD transduction | 610833/610832 |
| Anti p62/ SQSTM1 (C-terminus) | Guinea pig | Progen | GP62-C |
| Anti p62 | Mouse | Santa Cruz | sc-28359 |
| Anti IMMT/Mitofilin | Rabbit | Biomol | A305-023A-M |
| Anit MRPL45 (E-12) | Mouse | Santa Cruz | sc-515563 |
| Anti TOMM20 [EPR15581-54] | Rabbit | Abcam | ab186735 |
| Anti HADHA (E-8) | Mouse | Santa Cruz | sc-374497 |
| Anti HADHB (E-1) | Mouse | Santa Cruz | sc-271495 |
| Total OXPHOS Rodent WB Antibody Cocktail | Mouse | Abcam | ab110413 |
| Anti NIPSNAP1 | Rabbit | Abcam | ab67302 |
| Anti ODC (G-10) | Mouse | Santa Cruz | sc-390366 |
| AGO2 | Mouse | Abcam | ab57113 |
| SEC16A | Rabbit | Proteintech | 20025-1-AP |
| Normal IgG | Mouse | Santa Cruz | sc-2025 |

**Supplementary Table 6:** List of primers

| **Target** | **Sequence** |
| --- | --- |
| Hs mt-D Loop | Fwd: CTTCTGGCCACAGCACTTAAAC  Rev: GCTGGTGTTAGGGTTCTTTGTTTT |
| Hs mt-ND1 | Fwd: CCACCTCTAGCCTAGCCGTTTA  Rev: GGGTCATGATGGCAGGAGTAAT |
| Hs mt-ND6 | Fwd: CAAACAATGTTCAACCAGTAACCACTAC  Rev: ATATACTACAGCGATGGCTATTGAGGA |
| Hs mt-CYTB | Fwd: ATCACTCGAGACGTAAATTATGGCT  Rev: TGAACTAGGTCTGTCCCAATGTATG |
| Hs mt-CO1 | Fwd: GACGTAGACACACGAGCATATTTCA  Rev: AGGACATAGTGGAAGTGAGCTACAAC |
| Hs mt-ATP6 | Fwd: TAGCCATACACAACACTAAAGGACGA  Rev: GGGCATTTTTAATCTTAGAGCGAAA |
| Hs mt-RNR1 | Fwd: TAGAGGAGCCTGTTCTGTAATCGAT  Rev: CGACCCTTAAGTTTCATAAGGGCTA |
| Hs ß_2_M | Fwd: GCTGGGTAGCTCTAAACAATGTATTCA  Rev: CCATGTACTAACAAATGTCTAAAATGGT |
